# Predictive control of human pancreatic cell fate using a digital model of *in vitro* differentiation

**DOI:** 10.64898/2026.04.27.721124

**Authors:** Enrique E. Sanchez-Castro, Matthew Ishahak, Tai Le, Marlie M. Maestas, Diana C. Hernandez-Rincon, Noyonika Mukherjee, Kameron Bradley, James Lu, Sarah E. Gale, Jeffrey R. Millman

**Affiliations:** Division of Endocrinology, Metabolism and Lipid Research, Washington University School of Medicine; St. Louis, MO, USA; Department of Chemical, Environmental, and Materials Engineering, University of Miami; Coral Gables, FL, USA; Department of Biomedical Engineering, Washington University in St. Louis; St. Louis, MO, USA

**Author notes:** These authors contributed equally to this work (Enrique E. Sanchez-Castro, Matthew Ishahak).

**Keywords:** diabetes, embryonic stem cells, pluripotent stem cells, pancreatic islets, stem cell-derived islets, pancreatic differentiation, predictive modeling, transcription factors, single-cell sequencing, single-cell multiomic

## Abstract

The controlled generation of mature stem cell-derived islets (SC-islets) remains a barrier to scalable cell therapy for diabetes. Here, we develop a predictive digital model defining the cell-state-specific regulatory logic governing fate specification during human SC-islet differentiation. We integrate 400,603 cells from 9 original single-cell multiomic datasets and 52 public single-cell RNA-seq and ATAC-seq datasets across 4 cell lines and 7 differentiation protocols. This model resolves transcriptional and chromatin accessibility dynamics while enabling time-resolved inference and *in silico* perturbation of cell-state-specific gene regulatory networks. We identify regulators across trajectories from endoderm progenitors to pancreatic exocrine and endocrine lineages, nominating new candidate regulators. Among these candidates, we validate previously unreported roles for *STAT1* as an exocrine driver and *ZEB1* as a dynamic regulator of early endocrine specification and later off-target serotonergic islet cell fate. This work provides an experimentally supported predictive framework and an interactive resource comprising 1,116 simulations to prioritize transcription factors and intervention windows for refining SC-islet differentiation.

## INTRODUCTION

Recent clinical trials have demonstrated that stem cell-derived islets (SC-islets) can meaningfully restore glucose regulation in people living with type 1 diabetes^1–3^, supporting their promise as a scalable cell therapy for diabetes. However, current protocols still produce immature and heterogeneous SC-islets, containing incompletely specified pancreatic endocrine cells and off-target states^4^. These limitations indicate that SC-islet differentiation is not yet optimized to produce the cellular identity, maturity, and function desired for next-generation cell therapy.

Single-cell transcriptomic and chromatin accessibility studies have transformed the characterization of pancreatic differentiation by resolving cell identities, developmental intermediates, and molecular differences between *in vitro*-derived and primary islet cells^5–12^. Yet most single-cell atlases remain primarily descriptive, making it difficult to translate their insights into actionable interventions. A key unresolved challenge is predicting which regulatory factors should be perturbed, in which cell state, and at which developmental window to redirect fate specification^13^. Bridging this gap requires models that integrate temporal gene-expression and chromatin accessibility dynamics, infer cell-state-specific gene regulatory networks, and translate these networks into experimentally testable predictions.

Here, we introduce a predictive digital model of human *in vitro* pancreatic islet differentiation across multiple cell lines built from temporal single-cell epigenomic and transcriptomic data. This model transforms descriptive single-cell datasets into experimentally testable predictions of fate-specification regulators by integrating developmental trajectories with cell-state-specific gene regulatory networks (GRN). This approach recovers known lineage regulators and identifies previously unreported context-specific transcription factor (TF) activities, including roles for *STAT1* and *ZEB1*, which we validate *in vitro*. We also provide an interactive resource to explore predicted regulators and intervention windows across developmental stages and cell states (https://synbioelab.com/islettwin). Together, this work establishes a strategy for translating single-cell differentiation datasets into actionable approaches to refine SC-islet generation.

## RESULTS

### Multiomic mapping of SC-islet differentiation

Differentiation of human pluripotent stem cells into SC-islets relies on sequential exposure to small molecules and growth factors that mimic key *in vivo* developmental signals (**Fig. S1A-D**)^14^. This process yields functional organoids capable of dynamic, biphasic insulin secretion and amelioration of hyperglycemia in mice (**Fig. S1E-H**). To elucidate cell fate specification during the differentiation process, we jointly measured gene expression and chromatin accessibility using single-nucleus multiomic sequencing (snMulti-seq) across 9 time points, spanning 33 days, in 69,535 nuclei (**Fig. 1A and Fig. S2**). We resolved 13 developmental cell states based on assessment of canonical marker genes, chromatin accessibility at regions linked to identity genes, and activity of TF binding motifs over time (**Fig. 1B and Tables S1-S3**).

**Fig. 1.**
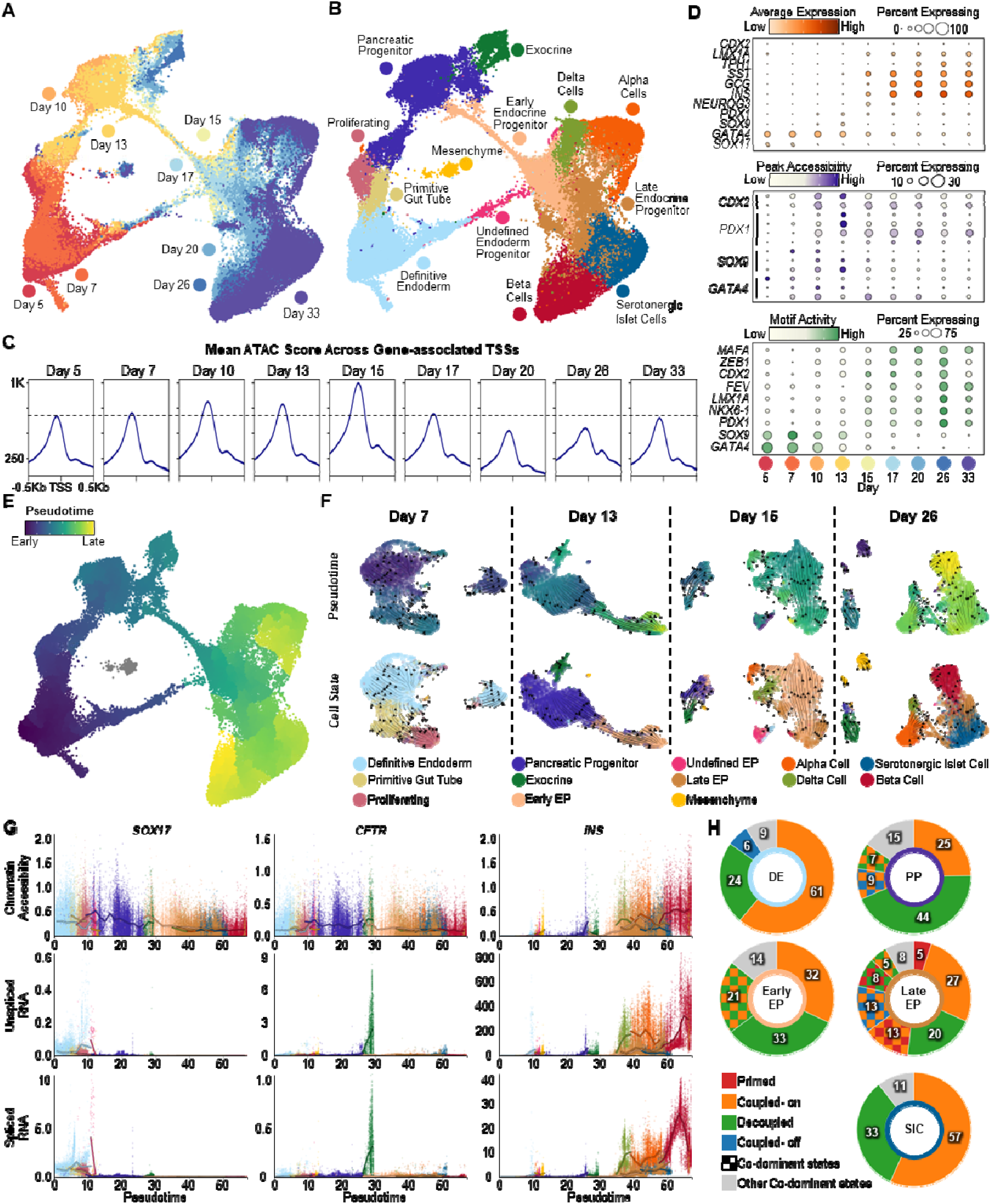
Multiomic mapping of transcriptional and epigenetic dynamics during SC-islet differentiation. **(A)** Multiomic UMAP of single-cell multiome data from SC-islet differentiation (69,535 cells), colored by differentiation day. **(B)** Multiomic UMAP colored by cell state annotations. **(C)** Mean normalized chromatin accessibility score around gene-associated transcription start sites (TSS) across differentiation time points. The dashed line indicates the score at day 5. **(D)** Dot plot summarizing gene expression (top), chromatin accessibility (middle), and transcription factor motif activity (bottom) for developmentally relevant markers across cell states throughout SC-islet differentiation. Dot size indicates the fraction of cells expressing each gene, and color indicates scaled average expression. **(E)** UMAP colored by gene expression-based pseudotime, highlighting progression through differentiation. **(F)** RNA velocity vectors overlaid on the UMAP for selected time points, with cells colored by pseudotime (top) or cell state (bottom). **(G)** Chromatin accessibility and RNA dynamics over pseudotime. For selected genes (SOX17 as a definitive endoderm marker, INS as a beta cell marker, and CFTR as an exocrine marker), scatter plots show pseudotime versus levels of normalized chromatin accessibility (top), unspliced pre-mRNA (middle), and spliced mature mRNA (bottom). Dots represent individual cells. Lines show the mean within each pseudotime bin for each cell state and are plotted only when ≥40 cells contribute. Dots and lines are colored by cell state. **(H)** Gene state composition across differentially expressed genes (DEGs), showing the percentage of DEGs classified as single-dominant or co-dominant across cells. A state is considered dominant when ≥25% of cells for a given gene fall into that state. “Other states” include dominant and co-dominant states that contribute <5%. DE, definitive endoderm; PP, pancreatic progenitor; EP, endocrine progenitor; SIC, serotonergic islet cells.

### Global chromatin accessibility peaks during endocrine induction as progenitor-state diversity expands

Pluripotent cells typically display broad chromatin accessibility that becomes restricted as lineage potential narrows^15,16^. To evaluate whether analogous trends occur during SC-islet differentiation, we quantified chromatin accessibility at transcription start sites (TSSs) across gene-associated DNA regions (**Fig. S3A**). We observed increasing accessibility around gene-associated TSSs until day 15, coincident with the start of endocrine lineage induction (**Fig. 1C**). This pattern suggests an increasingly open promoter landscape preceding endocrine induction, followed by progressive closing of chromatin as cells differentiate into more restricted endocrine lineages.

### Gene expression, chromatin accessibility, and motif activity show dynamic coupling and uncoupling during SC-islet differentiation

We next interrogated 1,500 differentially expressed genes (DEGs) to define how gene expression, chromatin accessibility, and transcription factor motif activity change during SC-islet differentiation. *GATA4* was highly expressed at the beginning of differentiation. This TF is required to prime the definitive endoderm (DE), establish the primitive gut tube (PGT), and activate the pancreatic gene program^17,18^. However, *GATA4*-associated chromatin accessibility and motif activity began to decline as cells exited the definitive endoderm state on day 7, followed by a delayed progressive decrease in *GATA4* expression starting on day 13 (**Fig. 1D** and **Fig. S3B**). *SOX9*, a TF required to establish early pancreatic progenitors (PP)^19,20^, high but decreasing motif activity from day 5, whereas peak chromatin accessibility and gene expression were observed from days 7 to 13 (**Fig. 1D** and **Fig. S3C**). *PDX1*, an essential regulator of early pancreatic lineage specification and endocrine differentiation^21,22^ showed strong correspondence among expression, peak accessibility, and motif activity starting from day 10, with increasing trends thereafter (**Fig. 1D** and **Fig. S3D).** Together, these examples show that transcriptional output, chromatin state, and inferred regulatory activity can be either temporally coupled or uncoupled during pancreatic lineage specification.

*NEUROG3*, a master regulator of endocrine induction^23,24^, showed transient, coordinated increases in expression and chromatin accessibility on days 15 and 17, consistent with progression toward endocrine lineages (**Fig. 1D** and **Fig. S3E**). Interestingly, expression of the pan-endocrine marker *CHGA* ^25,26^, preceded the *NEUROG3* activation and was detectable as early as day 13. More lineage-restricted endocrine markers, including *NKX6-1*, *INS*, *GCG*, and *SST*, emerged as early as day 15 and increased in both chromatin accessibility and expression thereafter (**Fig. 1D** and **Fig. S3F-G)**.

We also observed activation of markers associated with serotonergic programs, marked by expression and chromatin accessibility of *TPH1*, *LMX1A*, and *SLC18A1*. We defined this population as serotonergic islet cells (SIC) based on reports of developmental overlap between beta cells and serotonin-secreting neurons^27^, long-standing evidence of serotonin presence in pancreatic islets^28–31^, and limited expression of intestinal enterochromaffin markers, including *CDX2, TAC1,* and *SCT* (**Fig. 1D and Fig. S3H-I**)^32,33^.

### SC-islet differentiation shifts from unidirectional progression to progressive lineage divergence

We used our temporal multiomic data to reconstruct differentiation trajectories by combining transcriptional pseudotime ordering^34^ and stochastic velocity modeling^35^ (**Fig. 1E-F** and **Fig. S4**). Multiomic trajectory vectors aligned with the expected stagewise progression of SC-islet differentiation. DE cells at day 5 progressed largely unidirectionally toward the PGT state by day 7, followed by the pancreatic progenitor state by day 10. By day 13, trajectories began to diverge toward exocrine and early endocrine progenitor states, consistent with the bipotent potential of pancreatic progenitors during *in vivo* development. At later stages, vectors separated further as terminal endocrine states emerged.

### Gene-state analysis reveals context-specific chromatin-transcription regulatory modes

Mapping chromatin accessibility and transcriptional dynamics of DEGs over pseudotime revealed multiple regulatory modes during SC-islet differentiation (**Fig. 1G** and **Fig. S5**). Some genes followed the expected activation sequence in which chromatin accessibility increased before or alongside gene expression. This pattern was observed for several endocrine identity genes, including *INS*, *GCG,* and *SST*. However, other genes deviated from this canonical pattern. *SOX17*, a driver of definitive endoderm identity^36^, maintained chromatin accessibility throughout differentiation despite declining expression, indicating that transcriptional silencing can occur without immediate chromatin closure. *CFTR*, a ductal/exocrine marker, remained broadly accessible across multiple cell states but was transcribed specifically in exocrine cells, suggesting that open chromatin alone is insufficient for expression and that additional regulatory layers restrict lineage-inappropriate transcription. Together, these examples highlight that chromatin accessibility and transcription can be coordinated, delayed, or uncoupled depending on gene and cell-state context.

We characterized these patterns using four gene states^35^: primed (accessibility increases while expression remains low), coupled-on (accessibility and expression both increase), decoupled (accessibility and expression change in opposite directions), and coupled-off (accessibility and expression both decrease) (**Fig. S6A and Table S4**). Genes transitioned between dominant states across differentiation; therefore, we also defined co-dominant gene states to capture heterogeneous chromatin-transcription dynamics within a cell population (**Fig. S6B**). Most DEGs were classified as coupled-on, decoupled, or co-dominant for coupled-on and decoupled states, reflecting lineage programs that are actively induced, maintained, or transcriptionally silenced as trajectories progress. Canonical markers of pancreatic developmental stages, including *SOX17* in DE, *SOX9* in PP, and *NEUROG3* in early endocrine progenitors (EP), exhibited coupled-on dynamics (**Fig. S6C-E**). Other genes shifted from coupled-on to decoupled states, consistent with either stabilized expression or declining expression after maximal activation, as observed for *NOTCH1* in PP, *ESRRG* in early EP, and *CHGA* in late EP (**Fig. S6C-F**).

The largest proportion of primed genes was observed in the late EP cells (**Fig. S6D**), consistent with the previously observed global increase in chromatin accessibility prior to endocrine induction. Several terminal identity markers exhibited multistate behavior as these genes appeared and matured alongside the cell states, including *ERO1B* in beta cells, *GCG* in alpha cells, *SST* in delta cells, and *GLIS3* in exocrine cells (**Fig. S6G-J**). However, SICs were enriched for single dominant coupled-on or decoupled gene states, exemplified by *SLC18A1*, suggesting a more stable gene-state within this population (**Fig. S6K**).

### Constructing a predictive digital model of SC-islet differentiation

While our multiomic map provides insight into epigenetic and transcriptional dynamics during SC-islet differentiation, it does not capture the full breadth of trajectories that can arise across different protocols and cell lines. Therefore, we developed a framework to construct a more complete digital model of *in vitro* differentiation and enable robust downstream *in silico* analyses (**Fig. 2A**). We integrated 61 individual single-cell RNA sequencing (scRNA-seq), single-nucleus RNA sequencing (snRNA-seq), single-nucleus ATAC sequencing (snATAC-seq), and snMulti-seq datasets using our multiomic map as an anchor to generate a shared latent space and enable joint projection (**Fig. 2B-C and Table S5**). Annotation of the 400,603 cells within the digital model was based on protocol-dependent differentiation day and expression of canonical marker genes (**Fig. 2D and Figures S7-S13**). Based on the proportion of cell states observed across days, we identified six distinct stages that align with those reported by published protocols (**Fig. 2E**) ^37–43^. Stages 1 and 2 are dominated by cells in the DE and PGT states, respectively. In stage 3, we primarily observe cells in the PP state, followed by a shift to the early EP and late EP cell states in stage 4. Finally, stages 5 and 6 are characterized by increasing proportions of endocrine cell states. We reconstructed trajectory dynamics within the digital model by sampling random walks along a Markov chain constructed from multiomic transition probability matrices, which indicated alpha and beta cell states as the predominant endpoints (**Fig. 2F and Figure S14**). This framework provided a highly accurate model for predicting cell state transitions, with root mean square errors consistently below 0.05 for all cell states across differentiation stages (**Fig. 2G and Table S6**).

**Fig. 2.**
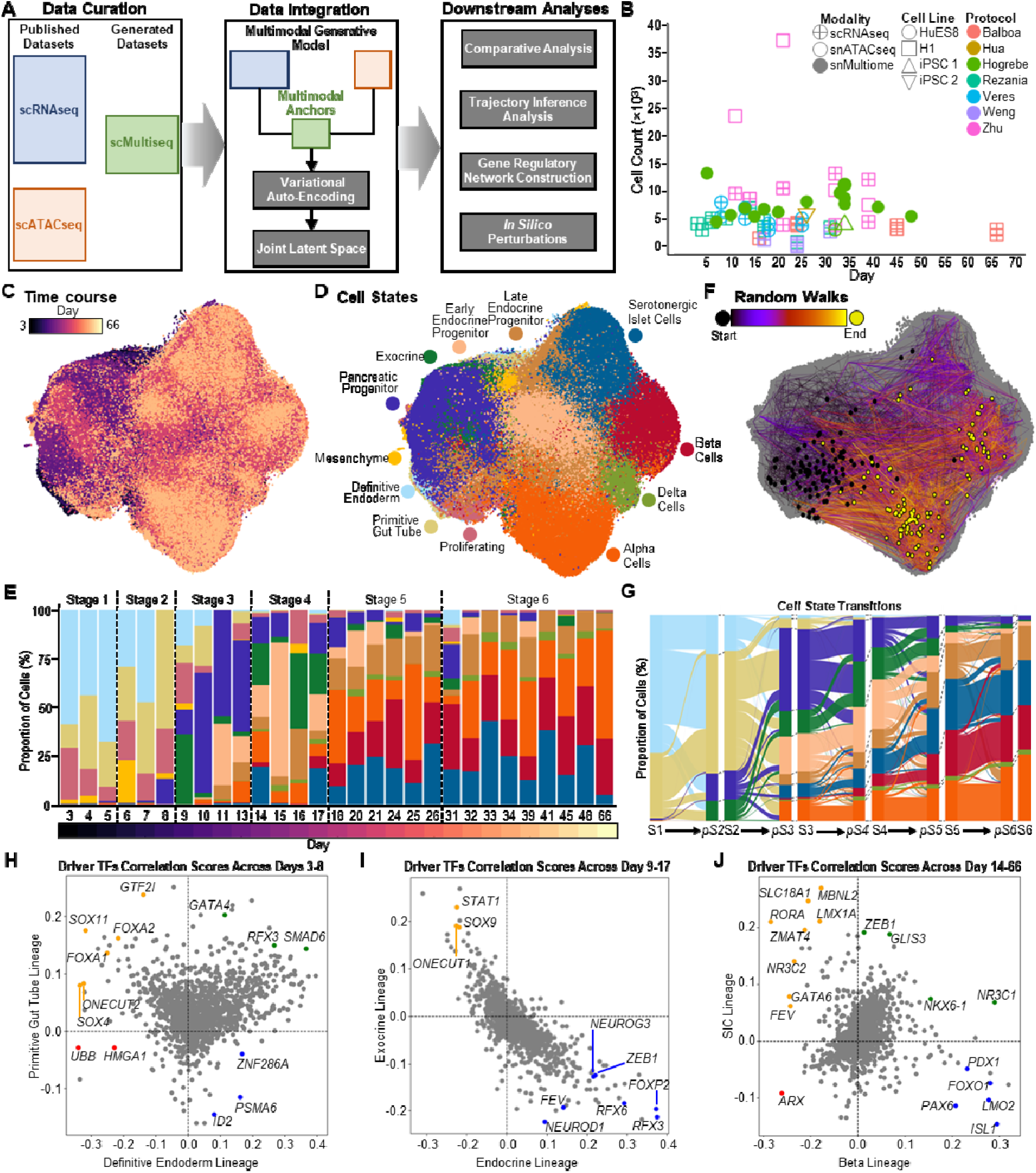
Constructing a multi-omic digital model that recovers known lineage regulators and nominates state-specific TF candidates. (**A**) Construction of the *in vitro* pancreatic islet differentiation digital model by curating public datasets and data newly generated in this study, applying a consistent preprocessing pipeline, and performing multimodal integration to enable downstream analyses (400,603 cells). (**B**) Metadata summary of the individual datasets used to construct the digital model. (**C**) UMAP of the *in vitro* pancreatic islet differentiation digital model colored by differentiation day. (**D**) UMAP of the *in vitro* pancreatic islet differentiation digital model colored by annotated cell state. (**E**) Proportion of annotated cell states acros differentiation days. Vertical dashed lines delineate differentiation stages. (**F**) Random walks visualized on the UMAP embedding. The start of each walk is marked by a black dot, consecutive stages are connected by edges, and yellow dots indicate walk endpoints. (**G**) Alluvial plots showing the proportion of cell states per stage and predicted transitions between consecutive stages. (**H**) Scatter plot summarizing driver TFs correlation scores between DE and PGT lineages from days 3-8. (**I**) Scatter plot summarizing driver TFs correlation scores between Endocrine and Exocrine lineages from days 9-17. (**J**) Scatter plot summarizing driver TFs correlation scores between Beta and SIC lineages from days 14-33.

### The digital model defines TFs correlated with cell-state-specific lineages

Using this digital model, we calculated the correlation of TFs with each major cell state across stages (**Table S7**). Developmentally related lineages, such as DE and PGT, shared substantial overlap in correlated transcription factors (**Fig. 2H**). By contrast, divergent branch points, including endocrine and exocrine specification, showed more lineage-restricted transcription factor correlations (**Fig. 2I)**. During early differentiation from day 3 to 8, *GATA4* and *FOXA2* were highly correlated with PGT specification (**Fig. 2H**)^18,44^. During pancreatic progenitor specification from day 9 to 17, transcription factors associated with exocrine development, including *ONECUT1* and *SOX9*, and endocrine development, including *NEUROG3*, *FEV*, *RFX3*, and *RFX6*, were highly correlated with their respective lineages (**Fig. 2I**)^45–49^. Notably, the model also identified previously unreported regulators at this bipotent specification stage, including *STAT1* for the exocrine lineage and *ZEB1* for the endocrine lineage **(Fig. 2I).**

Across the beta and SIC lineages from days 14 to 66, the model recovered established beta cell-associated regulators, including *PDX1*, *PAX6*, and *ISL1*, and SIC-associated regulators, including *SLC18A1*, *LMX1A*, and *FEV* ^22,32,50,51^ (**Fig. 2J**). *ZEB1*, *GLIS3, NR3C1*, and *NKX6-1* correlated with both lineages, suggesting partially shared regulatory programs (**Fig. 2J**). However, these correlations were lineage-biased with *ZEB1* and *GLIS3* more strongly associated with the serotonergic islet-cell lineage, whereas *NKX6-1* and *NR3C1* were more strongly associated with the beta lineage^52–54^. The model also identified regulators associated with other endocrine states, including *ARX* as selectively correlated with the alpha cell lineage, and *ISL1*, *PAX6*, and *MAFB* as shared across alpha and beta identities (**Table S7**) ^50,51,55–57^. Together, these results demonstrate that the integrated multiomic digital model recapitulates known lineage-regulatory programs associated with pancreatic development and provides insight into previously unexplored regulators of lineage specification.

### *In silico* transcription factor perturbations identify context-dependent regulators of early pancreatic progenitor specification

To further interrogate regulatory control during early SC-islet differentiation, we focused on the unidirectional transition from definitive endoderm to primitive gut tube and pancreatic progenitor states using 8,684 cells spanning days 5 to 13 from four independent differentiation protocols (**Fig. 3A–B** and **Fig. S15A-C**). Using gene expression-based pseudotime, we constructed developmental flow vectors that captured the trajectory of these early differentiation stages (**Fig. 3C**). We then inferred cell state-specific gene regulatory networks (GRNs) using an approach that incorporates both gene expression and chromatin accessibility data at regulatory genomic regions^58^. To summarize network structure, we ranked transcription factors by degree centrality, which quantifies the number of inferred regulatory connections and serves as a proxy for potential regulatory activity. Several highly connected transcription factors were shared across DE, PGT, and PP, including *TCF12*, *TCF7L2*, and *PRDM1* (**Fig. S15D**). By contrast, other transcription factors showed stage-dependent shifts in connectivity, with *FOXA1* decreasing and *CTCF* increasing as cells progressed from DE to early PP (**Fig. S15D** and **Table S8**).

**Fig. 3.**
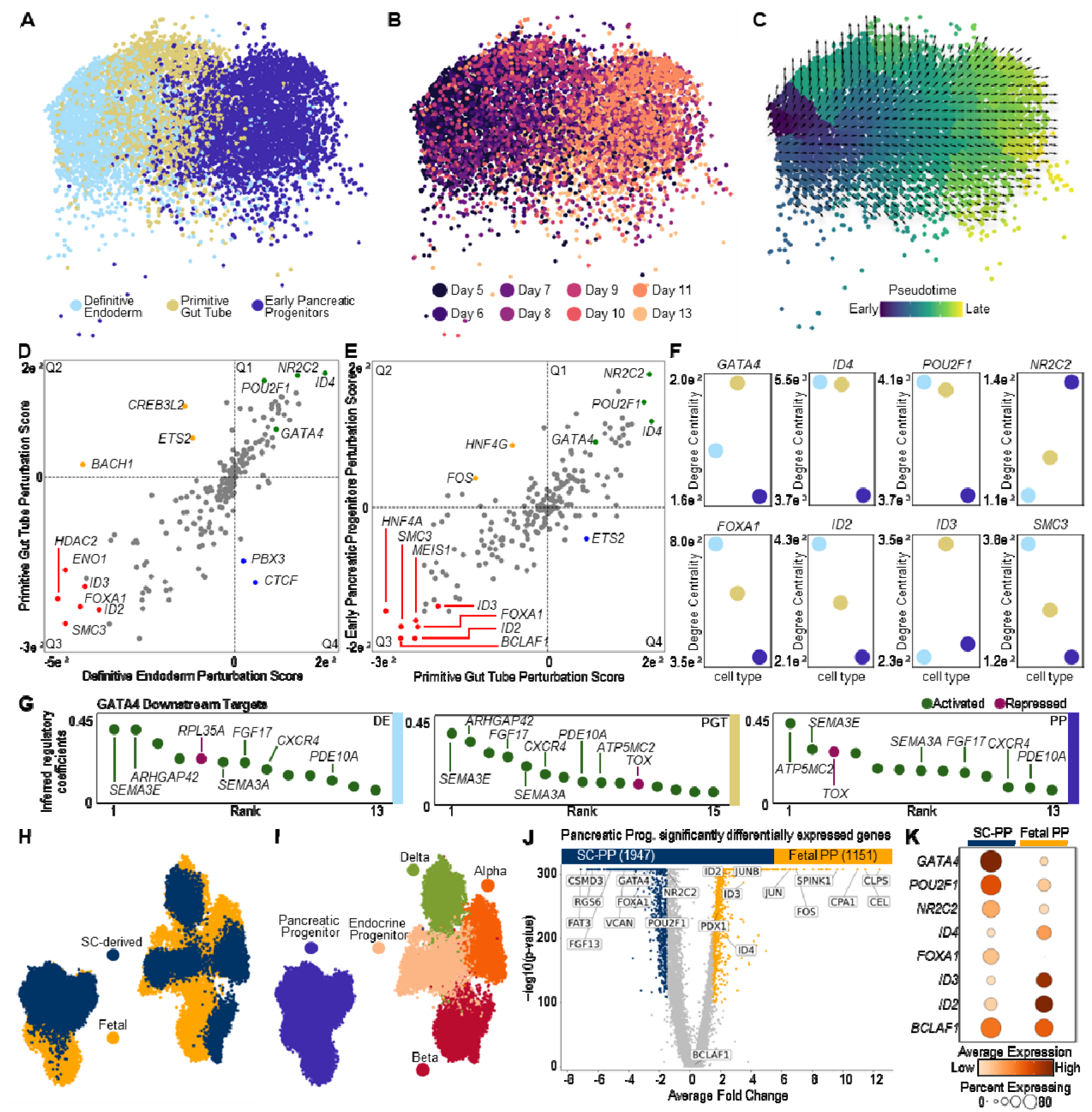
Early fate drivers during SC-islet differentiation diverge from fetal pancreatic development. (**A**) UMAP of the digital model focused on early pancreatic differentiation (8,684 cells), colored by cell state. (**B**) UMAP colored by differentiation day. (C) UMAP colored by gene expression-based pseudotime, with arrows indicating inferred developmental flow. (**D–E**) Scatter plots summarizing systematic TF knockout (KO) simulations. Each point represents a TF, positioned by mean perturbation score for definitive endoderm (x-axis) and primitive gut tube (y-axis) fates (D), or primitive gut tube (x-axis) and pancreatic progenitor (y-axis) fates (E). Quadrants indicate predicted effects: Q1, promote both fates; Q2, suppress the x-axis fate and promote the y-axis fate; Q3, suppress both fates; Q4, promote the x-axis fate and suppress the y-axis fate. Selected TFs are labeled and color-coded by quadrant (Q1 green, Q2 orange, Q3 red, Q4 blue). (**F**) Degree centrality of selected TFs with strong predicted perturbation effect across cell states (colors as in A). (**G**) Ranked inferred regulatory coefficients for *GATA4* targets in definitive endoderm (DE, left), primitive gut tube (PGT, middle), and pancreatic progenitors (PP, right), indicating predicted activation (green) or repression (purple). (**H**) UMAP of the integrated single-cell RNA-seq atlas (30,084 cells) combining in vivo human fetal pancreas and in vitro stem cell-derived pancreatic differentiation (Hogrebe protocol), colored by origin (fetal vs stem cell-derived). (**I**) UMAP colored by cell state. (**J**) Volcano plot of differentially expressed genes between stem cell-derived and fetal pancreatic progenitors. (**K**) Dot plot summarizing expression of selected TFs from KO simulations, comparing stem cell-derived and fetal pancreatic progenitors. Dot size indicates the fraction of cells expressing each gene, and color indicates scaled average expression.

Because GRN connectivity alone is insufficient to infer functional relevance, we simulated 222 individual transcription factor knockouts and quantified their predicted effects using a perturbation score (PS) calculated from the inner product of baseline developmental flow vectors and simulated KO vectors (**Fig. 3D-E** and **Table S9)**. KO of *ETS2*, *CREB3L2*, or *BACH1* was predicted to favor PGT specification while suppressing DE identity (**Fig. 3D**). Conversely, KO of *HNF4G* or *FOS* was predicted to promote early PP specification while suppressing PGT identity (**Fig. 3E**).

We next examined common predicted promoters and inhibitors of PGT and PP development. Predicted promoters included *ID4*, *POU2F1*, *NR2C2*, and *GATA4*, whereas predicted inhibitors included *ID2, ID3, SMC3,* and *FOXA1.* These TF showed distinct degree-centrality trends and scales, suggesting that their predicted effects were driven by the regulation of specific target genes rather than by overall network connectivity (**Fig. 3F** and **Fig. S15E**). Notably, KO of *GATA4* was predicted to promote progression toward both PGT and PP states, despite its established role in endoderm and pancreatic program induction^17,18^. We therefore examined inferred regulatory coefficients for putative *GATA4* targets across early cell states (**Fig. 3G** and **Table S10**). The model predicted that GATA4 positively regulated genes involved in tissue organization and pancreatic islet morphogenesis, including *SEMA3A*, *SEMA3E*, and *ARHGAP42*^59,60^. A positive coefficient was also observed for *FGF17*, a developmental signaling factor with mesoderm specification^61^, suggesting that high or persistent *GATA4* activity during *in vitro* differentiation may engage regulatory programs beyond those required for pancreatic lineage progression. Together, these observations suggest that early pancreatic specification depends on context-dependent transcription factor activity and that persistent activity of some developmental regulators, such as *GATA4*, may hinder rather than promote optimal *in vitro* progression.

### SC-islet differentiation shows persistent transcriptional overactivation relative to fetal *in vivo* **development**

To evaluate how closely *in vitro* differentiation recapitulates *in vivo* human pancreatic development, we integrated five independent scRNA-seq datasets spanning human development over post-conception weeks 4 to 19 with transcriptomic data from days 10 to 26 of our multiomic map (**Fig. 3H, Fig. S16A-C, and Table S11**). The integrated dataset comprised 30,084 cells representing PP, alpha, beta, and delta cell states (**Fig. 3I and Fig. S16D**). We then performed cell-state-matched DEG analysis between *in vivo* development and *in vitro* differentiation to identify 3,098 significant DEGs between stem cell-derived PPs (SC-PP) and fetal-derived PPs (**Fig. 3J and Table S12**), including several transcription factors predicted by our model to regulate *in vitro* PP specification. Notably, *GATA4*, *POU2F1*, and *NR2C2*, were overexpressed *in vitro*, while *ID2, ID3,* and *ID4* were overexpressed *in vivo* (**Fig. 3K**).

We also observed broad transcriptional overexpression across *in vitro* EP and terminal endocrine states compared with matched *in vivo* states, including a greater than six-fold increase in the number of DEGs in alpha and beta cells (**Fig. S17A-D**). This elevated transcriptional output is consistent with prior observations that SC-islets retain a more open chromatin landscape and persistent gene regulatory network activity than primary islets^11,62^. Interestingly, *GATA4* was recurrently upregulated across SC-derived states, with elevated expression across EP, alpha, beta, and delta lineages (**Fig. S17E**). These observations support the *in silico* prediction that sustained *GATA4* activity may constrain optimal *in vitro* progression and illustrate how the digital model can contextualize regulatory differences between SC-islet differentiation and human fetal pancreatic development.

### Branch-point modeling identifies regulators of exocrine-endocrine lineage choice

Next, we focused on a subset of 6,726 cells spanning days 11 to 24 from five independent *in vitro* differentiation protocols to explore the GRN landscape at the branch point between endocrine and exocrine lineages from the bipotent PP state (**Fig. 4A and Fig. S18A**). Within this subset, we resolved the divergent developmental flow from the PP state through two intermediate populations toward either the exocrine lineage or early EP lineage (**Fig. 4B-C).** While we observed decreased expression of *PDX1* along both the EP and exocrine lineages, the early EP intermediate population was characterized by increasing expression of *NEUROG3* and *CHGA*. On the other hand, the exocrine intermediate state had sustained expression of *SOX9* along with increasing expression of *CFTR* and *SPINK1*, which mark ductal and acinar lineages, respectively (**Fig. S18B-C**).

**Fig. 4.**
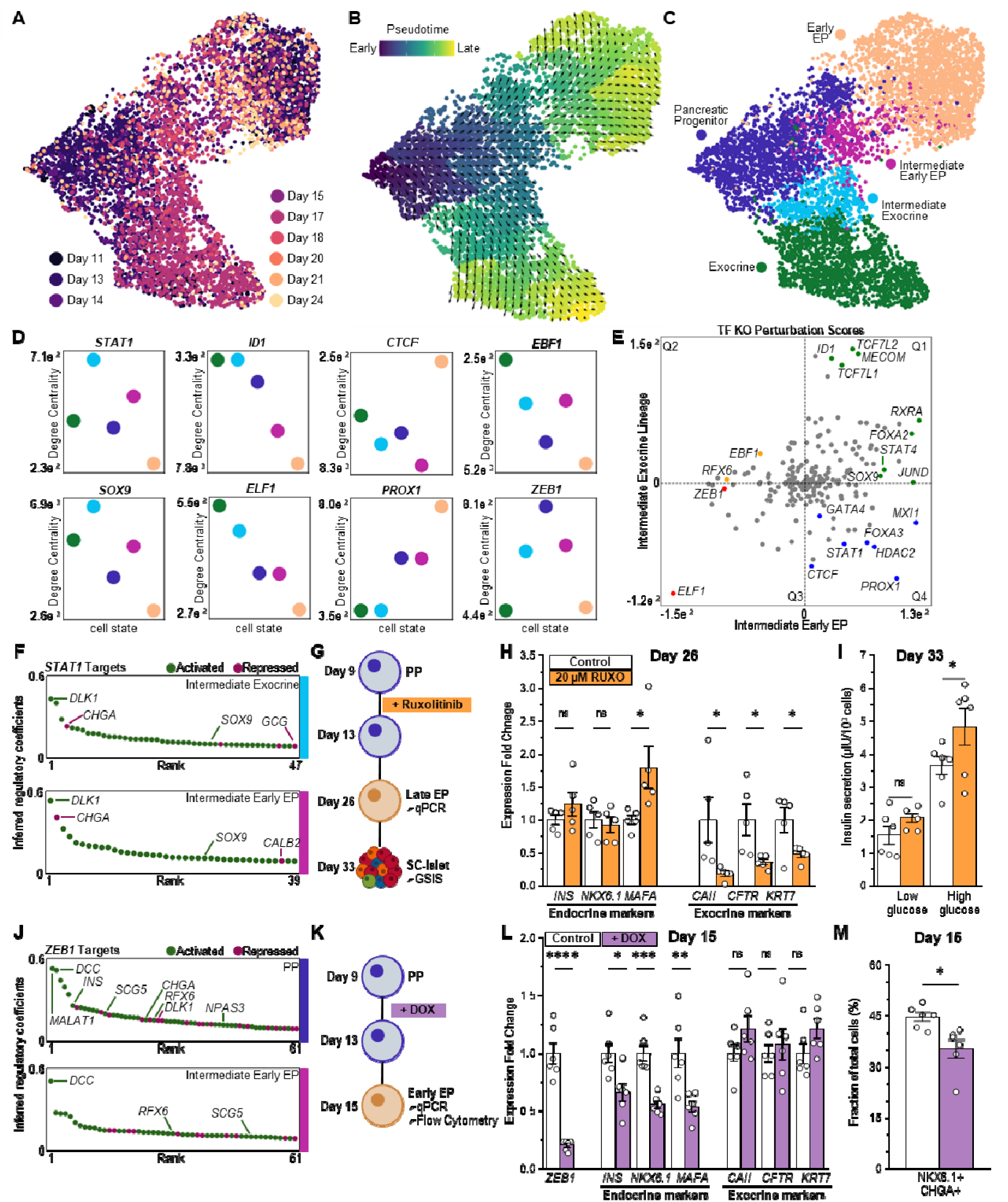
Exocrine-endocrine branch-point modeling reveals opposing roles for *STAT1* and *ZEB1* in lineage specification. (**A**) Subset UMAP of 6,726 cells from the digital model colored by differentiation day. (**B**) Subset UMAP colored by gene expression-based pseudotime, with arrows indicating inferred developmental flow. (**C**) Subset UMAP colored by cell state. (**D**) Degree centrality of selected TFs across cell states. (**E**) Scatter plot summarizing systematic TF knockout (KO) simulations. Each point represents a TF, positioned b mean perturbation score within the intermediate early EP GRN (x-axis) and intermediate exocrine GRN (y-axis). Quadrants indicate predicted effects: Q1, promote both fates; Q2, suppress the x-axis fate and promote the y-axis fate; Q3, suppress both fates; Q4, promote the x-axis fate and suppress the y-axis fate. Selected TFs are labeled and color-coded by quadrant (Q1 green, Q2 orange, Q3 red, Q4 blue). (**F**) Ranked inferred regulatory coefficients for *STAT1* targets in the intermediate exocrine GRN (top) and intermediate early EP GRN (bottom), indicating predicted activation (green) or repression (purple). (**G**) Schematic of ruxolitinib treatment (a JAK-STAT inhibitor) from days 9 to 13 during pancreatic progenitor specification. (**H)** qPCR at day 26 showing endocrine markers and exocrine markers. n = 5 biologically independent samples. (**I**) Static glucose-stimulated insulin secretion (GSIS) assay at day 33. Data are shown as mean ± s.e.m. relative to untreated control; n = 6 biologically independent samples. (**J**) Ranked inferred regulatory coefficients for *ZEB1* targets in the PP GRN (top) and intermediate early EP GRN (bottom), indicating predicted activation (green) or repression (purple). (**K**) Schematic of doxycycline (DOX)-inducible CRISPRi-mediated *ZEB1* knockdown from days 9-13 during pancreatic progenitor specification. (**L**) qPCR showing expression of *ZEB1*, endocrine markers, and exocrine markers. n = 6 biologically independent samples. (**M**) Flow cytometry quantification of NKX6.1+ and CHGA+ cells. Data are shown as mean ± s.e.m. relative to untreated control; n = 6 biologically independent samples. Statistical significance was assessed by unpaired two-sided t-test. * P < 0.05; ** P < 0.005; *** P < 0.0005; **** P < 0.00005. PP, pancreatic progenitor; EP, endocrine progenitor

We next inferred cell state-specific GRNs for each population and ranked TFs by degree centrality to summarize changes in network connectivity across the two lineages (**Table S13**). *HDAC2*, which encodes a protein responsible for transcriptional repression and chromatin remodeling^63^, was highly connected across both lineage trajectories (**Fig. S18D**). This finding aligns with our previous observation that chromatin accessibility across TSSs begins to decrease genome-wide after day 15 of *in vitro* differentiation (**Fig 1D**). We also observed other TFs that had state-specific connectivity patterns (**Fig. 4D**). For example, *STAT1* and *SOX9* connectivity increased in both intermediate populations, but this increased connectivity was maintained only along the exocrine lineage. Other TFs showed similar lineage-specific connectivity changes, with *ID1*, *EBF1,* and *ELF1* increasing along the exocrine trajectory, whereas *CTCF* and *PROX1* increased connectivity along the early EP trajectory. By contrast, TFs such as *ZEB1* showed maximal connectivity in PP and then decreased connectivity along both lineages.

To prioritize TFs that may regulate lineage specification at this branch point, we calculated PS from systematic *in silico* TF KO simulations across 226 TFs with inferred regulatory connections in the intermediate population GRNs (**Fig. 4E** and **Table S14**). Many KOs were predicted to promote both intermediate fates, including KO of *ID1*, *TCF7L2*, *MECOM*, *FOXA2*, or *RXRA* (**Fig. 4E**). In contrast, the KO of *STAT1*, *HDAC2*, or *PROX1* was predicted to specifically promote progression toward the intermediate early EP lineage while suppressing the intermediate exocrine lineage (**Fig. S18E**). Additionally, KO of *RFX6* or *ZEB1*, which are both regulators implicated in pancreatic endocrine development and function^48,64^, were predicted to strongly impede progression toward the intermediate early EP state while minimally affecting the intermediate exocrine lineage. Notably, several TFs that were found to be upregulated relative to *in vivo* development were also predicted to affect early EP differentiation upon simulated KO, including *STAT1*, *RFX6*, and *ZEB1* (**Fig. S9E and S18E**).

### *STAT1* and *ZEB1* regulate opposing fates during the exocrine-endocrine lineage specification

We examined the inferred downstream targets of *STAT1*, since it has not been previously reported as a regulator of pancreatic development, to uncover the potential regulatory mechanism underlying the *in silico* observations (**Fig. 4F and Table S15**). We found that *STAT1* was predicted to positively regulate genes including *DLK1* and *SOX9* while repressing *CHGA* in both intermediate states. To test the *in silico* observation, we inhibited *STAT1* transcription using a doxycycline-inducible CRISPR interference (CRISPRi) system during days 9 to 13 of *in vitro* differentiation (**Fig. S18F**). Activation of the CRISPRi construct resulted in a significant decrease in *STAT1* expression on day 13 which was reversed by day 26 (**Fig. S18G-H**). On day 26, we also found that the temporary knockdown of *STAT1* did not alter expression of beta cell markers (*INS*, *NKX6-1*, *MAFA*) while significantly reducing expression of exocrine markers, such as *CAII* and *CFTR* (**Fig. S18I**). To further validate the role of *STAT1* in exocrine lineage specification, we pharmacologically inhibited JAK–STAT signaling using ruxolitinib (RUXO) over the same timeframe and again profiled cells on day 26 (**Fig. 4G**). We found that RUXO treatment preserved or improved expression of beta cell markers, while significantly reducing expression of exocrine markers (**Fig. 4H**). Functional assessment on day 33 demonstrated significantly increased insulin secretion in response to high glucose stimulation following RUXO treatment (**Fig. 4I**).

We also examined *ZEB1*, which has been reported as a regulator of pancreatic development in mice^64^, but has not been studied in SC-islets. Interestingly, many endocrine-associated genes, including *INS*, *DCC*, *SCG5*, *RFX6*, *CHGA*, and *NPAS3*, were among the inferred downstream targets of *ZEB1* within the PP GRN (**Fig. 4J**, top). However, many of these genes did not show up as downstream targets in the intermediate early EP GRN (**Fig. 4J**, bottom), suggesting a role as an early regulator of endocrine specification. We utilized our CRISPRi system to test the *in silico* prediction that *ZEB1* knockdown would hinder endocrine specification (**Fig. 4K**). On day 15, we found that *ZEB1* knockdown significantly reduced expression of endocrine markers, while exerting no significant effect on exocrine markers (**Fig. 4L**). Concordantly, protein-level quantification showed a significant reduction in the fraction of NKX6-1^+^/ CHGA^+^ cells (**Fig. 4M**). Together, these results support the *in silico* predictions that *STAT1* and *ZEB1* play a role in exocrine and endocrine specification, respectively, and highlight the utility of our digital model for identifying regulators of cell fate during *in vitro* differentiation.

### Lineage-specific GRN modeling of regulators during alpha and delta cell specification

Attempts to control endocrine lineage specification during SC-islet differentiation have had limited success^65–68^. Therefore, we sought to characterize the GRNs driving alpha and delta cell lineages. To interrogate regulatory control along alpha lineage specification during *in vitro* differentiation, we generated a subset consisting of 16,122 cells from days 10 to 21 (**Fig. S19A-E**). Consistent with previous reports of multiple alpha cell lineage trajectories^10^, we observed one trajectory progressing from the PP through an early alpha intermediate state and a second trajectory progressing through the early EP state (**Fig. S19F**). Inferred GRNs for each population revealed general transcriptional regulators associated with human development, such as *TCF12* and *JUND*, to be highly connected across all lineage states (**Fig. S19G and Table S16**). To identify TFs that may regulate these two alpha cell lineage trajectories, we performed *in silico* TF KO simulations for all 227 TFs within this subset. Expectedly, we found that KO of canonical endocrine and alpha-associated TFs^55–57,69^, such as *ARX*, *MAFB*, *PAX6*, and *NR3C1*, blocked progression along both alpha trajectories (**Fig. S19H-J and Table S17**).

We also investigated the regulation of delta cell lineage specification using a subset of 7,583 cells spanning days 11 to 21 (**Fig. S20A-C**). Within this subset, we observed an intermediate delta state characterized by lower expression of identity markers, such as *SST* and *HHEX*, and slightly higher expression of *NEUROG3* (**Fig. S20D-E**). The developmental flow profile revealed a unidirectional trajectory from PP through EP states into the intermediate delta and delta states (**Fig. S20F**). Inferred GRNs again revealed TFs associated with broad transcriptional regulation, including *TCF12*, *JUND*, and *HDAC2*, were highly connected across cell states (**Fig. S20G and Table S18**). *In silico* TF KO simulations of the 227 TF in this subset identified *PBX3*, *FOXO1*, *NR2C2*, and *NF1* as regulators of delta lineage progression (**Fig. S20H-J and Table S19**). These factors have been reported to play diverse roles in pancreatic development, endocrine function, and somatostatinoma^70–73^. While we did not delve into specific regulators of the alpha and delta lineages, we provide the inferred regulatory coefficients for downstream TF targets within the cell-state-specific GRNs (**Tables S20-S21**) and encourage the use of our online tool (https://www.synbioelab.com/islettwin) to facilitate future mechanistic studies.

### *ZEB1* shifts from supporting endocrine progenitor specification to promoting SIC fate at beta/SIC bifurcation

Previous reports suggest that the off-target SIC lineage is closely related to beta cell specification; therefore, we interrogated these trajectories using a subset of 6,922 cells from days 15 to 21 (**Fig. 5A and Fig. S21A**). Within this subset, we defined early EP, late EP, beta, and SIC states based on expression of canonical identity markers (**Fig. 5B and Fig. S21B-C**). Interestingly, we observed a hybrid population expressing markers of both beta and SIC lineages, and the inferred developmental flow vectors revealed a slight bifurcation toward either lineage, suggesting subtle differences between the two trajectories (**Fig. 5B-C**). We next inferred cell state-specific GRNs and ranked transcription factors by degree centrality to summarize changes in network connectivity (**Fig. S21D and Table S22**). Several highly connected regulators were shared across cell states, including *TCF12*, *HDAC2*, and *BCLAF1*. Other genes demonstrated lineage-associated shifts in connectivity as cells progressed toward either SIC or beta cell lineages. For example, *ZEB1* ranked tenth in degree centrality within the late EP GRN and jumped to the top five in the SIC and hybrid GRNs. In contrast, *RXRA* and *MAFB* only ranked among the top 20 connected TFs in the beta GRN. Degree centrality scores across cell states for eight TFs further revealed divergent TF activity between beta and SIC lineages (**Fig. 5D**). *HNF4A* and *ESRRG* were highly connected within the early EP GRN. Connectivity of *ZEB1* and *LMX1A* peaked in the SIC and hybrid GRNs. *PAX4* and *FEV* showed high connectivity in both the SIC and beta GRNs, while *KLF12* and *TCF7L2* peaked in the hybrid GRN.

**Fig. 5.**
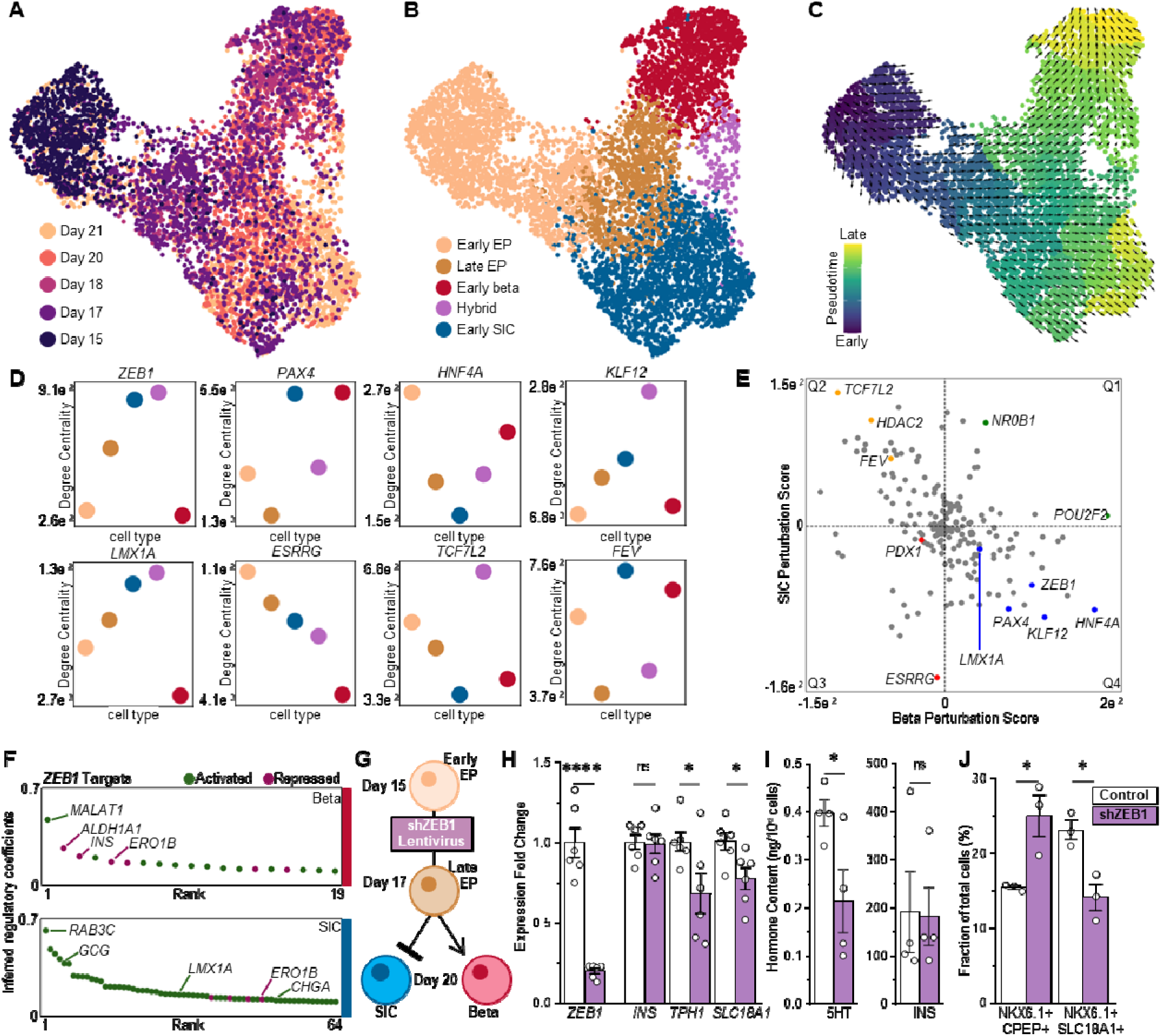
Beta-SIC branch-point modeling reveals a stage-dependent shift in ZEB1 activity that promotes SIC fate. (**A**) UMAP of the digital model focused on the Beta and SIC branching point (6,922 cells), colored by cell state. (**B**) UMAP colored by differentiation day. (**C**) UMAP colored by gene expression-based pseudotime, with arrows indicating inferred developmental flow. (**D**) Degree centrality of selected TFs with strong predicted perturbation effects across cell states. (**E**) Scatter plot summarizing systematic TF knockout (KO) simulations. Each point represents a TF, positioned by mean perturbation score for Beta (x-axis) and SIC (y-axis) intermediate fates. Quadrants indicate predicted effects: Q1, promote both fates; Q2, suppress the x-axis fate and promote the y-axis fate; Q3, suppress both fates; Q4, promote the x-axis fate and suppress the y-axis fate. Selected TFs are labeled and color-coded by quadrant (Q1 green, Q2 orange, Q3 red, Q4 blue). (**F**) Ranked inferred regulatory coefficients for *ZEB1* targets in Beta (top) and SIC (bottom), indicating predicted activation (green) or repression (purple). (**G**) Schematic representation of hypothesis that knockdown of *ZEB1* via shRNA on days 15-17 will inhibit SIC lineage progression without hurting beta cell specification. (**H**) qPCR showing expression of *ZEB1, INS, TPH1*, and *SLC18A1* on day 20. n = 6 biologically independent samples. (**I**) Hormone content on day 20 for serotonin (5HT, left) and insulin (INS, right). n = 4 biologically independent samples; Data are shown as mean ± s.e.m. (**J**) Flow cytometry quantification of NKX6-1+/C-peptide+ (Beta cells) and NKX6-1+/SLC18A1+ (SIC cells), shown as mean ± s.e.m. relative to GFP shRNA control. n = 3 biologically independent samples. Statistical significance was assessed by unpaired two-sided t-test. * P < 0.05; ** P < 0.005. EP, endocrine progenitor; SIC, serotonergic islet cells.

We then performed systematic *in silico* TF KO simulations across 214 TFs with inferred regulatory connections to identify candidate regulators of the fate decisions between beta and SIC states (**Fig. 5E and Tables S23-S24**). KO of *NR0B1* was predicted to promote both SIC and beta fates, whereas KO of *PDX1* was predicted to suppress both lineages. KO of *TCF7L2*, *HDAC2*, or *FEV* was predicted to promote SIC specification while suppressing the beta lineage. Conversely, KO of *PAX4*, *LMX1A*, or *ZEB1* was predicted to promote beta specification while suppressing the SIC lineage. Representative KO vector-field projections and TF expression patterns further contextualized predicted effects within the relevant cell populations (**Fig. S21E**). Notably, increased expression of *ZEB1* was observed along the SIC lineage compared to both late EP and beta cell states. Examination of the downstream targets of *ZEB1* within the beta and SIC GRNs further highlighted cell-state-specific activity. Within the beta GRN, ZEB1 was inferred to repress a subset of beta cell identity genes, including *INS* and *ERO1B* **(Fig. 5F, top)**. In the SIC GRN, *ZEB1* retained a repressive effect on *ERO1B* with an activator effect on *CHGA* and *LMX1A* **(Fig. 5F, bottom**). This illustrates the temporal and cell-state dependence of TF activity: *ZEB1* not only changed its inferred targets across differentiation but also shifted from activating *INS* in the PP state to repressing it in beta cells.

To test our *in silico* observations, we delivered a short hairpin RNA (shRNA) construct via lentivirus to induce *ZEB1* knockdown on days 15 to 17 (**Fig. 5G**). On day 20, we found that *ZEB1*, *TPH1*, and *SLC18A1* expression were significantly reduced, while *INS* expression was not affected (**Fig. 5H**). This corresponded with a significant decrease in serotonin content and no change in insulin content (**Fig. 5I**). Finally, assessment of cell identity markers by flow cytometry revealed a significant increase in NKX6-1⁺/C-peptide⁺ beta cells and a significant decrease in NKX6-1⁺/SLC18A1⁺ SIC proportions (**Fig. 5J**). These results indicate a dynamic role for *ZEB1*, where early activity is necessary for endocrine lineage specification, but continued activity disrupts beta cell specification.

## DISCUSSION

Understanding how cell fate is regulated during *in vitro* differentiation is essential for improving SC-islet cell replacement therapies for diabetes. Although current protocols generate functional endocrine cells, precise control of endocrine specification remains limited, resulting in off-target populations^65–68^. Here, we present a predictive digital model of SC-islet differentiation across multiple cell lines that integrates transcriptional and chromatin accessibility dynamics to reconstruct developmental trajectories, correlate TF drivers with major cell states, infer cell-state-specific GRNs, and investigate candidate regulators via *in silico* TF perturbations. This framework moves beyond a descriptive atlas of *in vitro* differentiation by providing an interactive model of regulatory control across DE-to-PP progression, exocrine-endocrine bifurcation, and terminal endocrine lineage decisions.

A major insight from our multiomic assessment is that SC-islet differentiation does not follow the expected trend in which chromatin accessibility progressively decreases as cells transition from immature progenitors to terminal endocrine states^16,74^. Instead, chromatin accessibility across TSSs increased until the endocrine induction window, coinciding with divergent developmental flow and enrichment of primed gene states in EP cells. These findings suggest that chromatin accessibility reflects the differentiation potential of progenitor populations and can transiently increase before multi-lineage specification events. We also found that gene expression, chromatin accessibility, and TF motif activity can be temporally coupled or uncoupled depending on gene and cell-state context, consistent with our previous comparative analyses^11,62^. While some genes followed the expected pattern in which chromatin opening preceded or accompanied transcriptional activation, others remained accessible despite transcriptional silencing or were expressed only in specific lineages despite broader accessibility. Thus, chromatin accessibility alone is insufficient to infer transcriptional output, and vice versa^75^, highlighting the value of paired snMulti-seq measurements for building integrated multiomic GRNs^76,77^.

The digital model also enabled comparison between *in vitro* differentiation and human pancreatic development *in vivo*. Although SC-islet differentiation captured major developmental transitions, it did not fully recapitulate the extent or timing of transcriptional repression observed during fetal pancreatic development. *In vitro*-derived pancreatic progenitor and endocrine states showed broad transcriptional overexpression relative to matched fetal states, including recurrent overexpression of *GATA4* across multiple SC-derived lineages. This finding contextualizes the *in silico* prediction that sustained high *GATA4* activity may constrain early pancreatic specification, despite the established requirement for *GATA4* to develop the pancreas^17,18,78^. More broadly, our observations indicate that *in vitro* differentiation is not an accelerated version of fetal development, but a related process with distinct regulatory mechanisms and lineage-state constraints.

Simulated TF perturbations further demonstrate the importance of context-specific regulatory activity. The model recovered expected roles for established pancreatic regulators, including *FOXA2*, *SOX9*, *NEUROG3*, *RFX6*, *ARX*, *PDX1*, and *NKX6-1*,^19,22,44,49,52,57,79^ supporting the biological validity of the inferred GRNs. At the exocrine-endocrine branch point, *in silico* TF KO simulations suggested roles for *STAT1* and *ZEB1* as lineage regulators with opposing lineage effects. *In vitro* studies supported these predictions with transient *STAT1* inhibition resulting in reduced exocrine marker expression and improved glucose-stimulated insulin secretion, whereas early *ZEB1* knockdown impaired endocrine specification. During downstream beta cell specification, however, *in silico* simulations suggested *ZEB1* activity promoted the off-target SIC fate, which was validated by *in vitro ZEB1* knockdown resulting in reduced SIC markers and serotonin content while increasing the proportion of beta cells. These results illustrate that TF function is not fixed, but depends on developmental timing, cell state, and target-gene context.

Overall, this work establishes a predictive framework for translating single-cell multiomic data into experimentally testable strategies to improve *in vitro* differentiation efficiency. By integrating temporal multiomic measurements with GRN inference and *in silico* perturbation, the model identifies conserved and context-specific regulators of pancreatic lineage specification. More broadly, this study illustrates how predictive models can build on single-cell characterization of stem cell differentiation by connecting cell-state maps to testable regulatory hypotheses, rational strategies to refine cell identity, reduced off-target fate specification, and improved future cell replacement therapies for diabetes.

## RESOURCE AVAILABILITY

### Lead contacts

• Requests for further information and resources should be directed to and will be fulfilled by the lead contacts, Jeffrey R. Millman and Matthew Ishahak.

### Materials availability

- This study did not generate new unique reagents.

### Data and code availability

- The snMulti-seq data generated in this study have been deposited in the Gene Expression Omnibus (GEO) database.
- Code was developed and executed using a custom containerized Linux environment consisting of JupyterLab v4.0.11, R v4.4.1, and Python v3.10.13 available on Docker Hub (https://hub.docker.com/r/ishahakm/leary).
- The scripts used for analysis are available on GitHub (https://github.com/ishahakm/scisletdigitaltwin).
- All other data are available in the main text or supplementary materials.
- Any additional information required to reanalyze the data reported in this paper is available from the lead contact upon request.

## ACKNOWLEDGMENTS

We would like to thank the McDonnell Genome Institute (MGI) at Washington University School of Medicine and the Research Infrastructure Services (RIS) at Washington University in St. Louis for the generation and storage of sequencing data, respectively. We would also like to thank Erika Brown (Washington University) for her feedback on the manuscript. We also thank Nathaniel Hogrebe, Punn Augsornworawat, Daniel Veronese-Paniagua, Alice Smagorinsky, Maddy Goedegebuure, Chris Walter, and Nathaniel Hogrebe for additional support throughout this project.

Washington University occupies the ancestral, traditional, and contemporary lands of the Osage Nation, Missouria, Illinois Confederacy and many other tribes as the custodians of the land where we reside, occupy, and call home. We recognize their sovereignty was never ceded after unjust removal and encourage your own research on tribal removal, tribal sovereignty and the history of the land you reside. We promote the inclusion of tribal history and the incorporation of contemporary thoughts and actions into your work. In offering this land acknowledgement, we affirm and support Tribal sovereignty, history and experiences by elders past, present, and seven generations yet to come through their continued connection to this land.

We also thank our funding sources:

National Institutes of Health grant UG3DK142188 (JRM)

National Institutes of Health grant T32DK007120 (MI)

National Institutes of Health subawards from U24DK104162

HIRN Catalyst Award 61294.2006834.669310 (JRM)

HIRN Data Scholars Program 66220.2013345.669302 (MI)

Breakthrough T1D grant 3-SRA-2023-1295-S-B (JRM)

Edward J. Mallinckrodt Foundation (JRM)

Beatson Foundation grant 2022-001 (JRM)

Anita Palmer Corbin Trust (JRM)

Rita Levi-Montalcini Postdoctoral Fellowship in Regenerative Medicine (MI and NM)

## AUTHOR CONTRIBUTIONS

Conceptualization: EESC, MI, JRM

Data curation: MI

Formal analysis: EESC, MI

Funding acquisition: MI, JRM

Investigation: EESC, MI, TL, MM, DCHR, KB, NM, JL

Methodology: EESC, MI, JRM

Project administration: MI, JRM

Resources: EESC, MI, SEG, JRM

Software: EESC, MI

Supervision: MI, JRM

Validation: EESC, MI, TL

Visualization: EESC, MI

Writing – original draft: EESC, MI, JRM

Writing – review & editing: EESC, MI, TL, MM, KB, NM, JL, SEG, JRM

## DECLARATION OF GENERATIVE AI AND AI-ASSISTED TECHNOLOGIES IN THE WRITING PROCESS

During the preparation of this work, the authors used ChatGPT, powered by GPT-5.5 Thinking, to assist with improving the readability, clarity, and language of the manuscript. The authors reviewed and edited the content as needed and take full responsibility for the content of the publication.

## SUPPLEMENTAL INFORMATION

Document S1. Supplemental Tables S1–S25

Document S2. Supplementary Figures S1-S21

## FIGURE TITLES AND LEGENDS

**Fig. S1 *In vitro* generation and characterization of stem cell-derived islets.**

(**A**) Overview of 6-stage in vitro differentiation protocol used to generate SC-islets. (**B**) Representative brightfield images indicating morphological changes during in vitro differentiation at low (top) and high (bottom) magnification. Scale Bars = 100 µm. (**C**) Immunofluorescent imaging of temporal protein expression during in vitro generation of SC-islets. Scale Bar = 100 µm. (**D**) Temporal expression of canonical marker genes associated with developmental states. n=3 biological replicates. (**E**) Static glucose stimulated insulin secretion (sGSIS) of SC-islets on Day 33. Simulation index calculated as insulin concentration at high glucose (20mM) divided by insulin concentration at low glucose (2mM). n=20 biological replicates. (**F**) Dynamic glucose stimulated insulin secretion of SC-islets on Day 33 demonstrating biphasic insulin secretion. n=5 biological replicates; Data are shown as mean ± s.e.m. (**G**) Fasting blood glucose of streptozotocin-treated mice after transplantation of approximately 2.29 ± 0.32 × 106 cells per mouse. Green region indicates normoglycemic range. n=8 mice per condition; Data are shown as mean ± s.e.m. (H) Glucose tolerance test 3-months after SC-islet transplantation. n=8 mice per condition; Data are shown as mean ± s.e.m.

**Fig. S2 Quality control, integration, and annotation of integrated multi-omic atlas.**

(**A**) Violin plots of quality control metrics for single-nucleus multiome sequencing grouped by time point, including RNA counts per nucleus (nCount_RNA), ATAC fragment counts per nucleus (nCount_ATAC), transcription start site (TSS) enrichment, and nucleosome signal (ratio of mononucleosomal to nucleosome-free fragments). (**B**) UMAPs generated from RNA data only using different integration methods colored by differentiation day. (**C**) Integration benchmarking summary of method performance across batch-correction and biological conservation metrics. (**D**) UMAPs generated from only RNA data (top) or only ATAC data (bottom) colored by differentiation day. (**E**) Joint weighted nearest neighbor UMAP colored by Louvain clusters. (**F**) Bar plot showing the proportion of cell states identified at each day of SC-islet differentiation.

**Fig. S3 Characterization of chromatin accessibility during SC-islet differentiation.**

(**A**) Tornado plots showing genome-wide chromatin accessibility profiles for peak sets across differentiation time points. TSS: Transcription Start Site. (**B**) Coverage plots across differentiation days showing chromatin accessibility at *GATA4* gene locus. (**C**) Coverage plots across differentiation days showing chromatin accessibility at *SOX9* gene locus. (**D**) Coverage plots across differentiation days showing chromatin accessibility at *PDX1* gene locus. (**E**) Coverage plots across differentiation days showing chromatin accessibility at *NEUROG3* gene locus. (**F**) Dot plot summarizing gene expression across differentiation days for TFs associated with endocrine cell states. (**G**) Coverage plots across differentiation days showing chromatin accessibility at gene loci for pancreatic endocrine hormones. (**H**) Dot plot showing lack of gene expression of intestinal enterochromaffin markers. (**I**) Coverage plots across differentiation days showing chromatin accessibility at gene loci for intestinal enterochromaffin markers.

**Fig. S4 Multiomic velocity during different days of SC-islet differentiation.**

RNA velocity vectors are overlaid on the UMAP for individual time points, with cells colored by pseudotime (left) or cell state (right).

**Fig. S5 Chromatin accessibility and RNA dynamics over pseudotime for selected genes.**

Scatter plots show pseudotime versus levels of normalized chromatin accessibility (left), unspliced pre-mRNA (middle), and spliced mature mRNA (right). Seven representative genes are shown: *OTX2* (**A**), *GATA4* (**B**), *CHGA* (**C**), *NKX6-1* (**D**), *GCG* (**E**), *SST* (**F**), and *SLC18A1* (**G**). Dots represent individual cells. Lines show the mean within each pseudotime bin for each cell state and are plotted only when ≥40 cells contribute. Dots and lines are colored by cell state.

**Fig. S6 Gene State distributions and dynamics.**

(**A**) Schematic illustrating four gene states. Gene expression and chromatin accessibility are coupled in the coupled-on (orange) and coupled-off (blue) states and decoupled in the primed (red) and decoupled (green) states (adapted from ref. 34). (**B**) Percentage of differentially expressed genes (DEGs) classified with single-dominant or co-dominant gene states across cell states. A gene state is considered dominant when ≥25% of cells for a given gene fall into that gene state. “Other states” groups single dominant and co-dominant gene states contributing <5%. (**C-K**) Scatter plots show gene time versus levels of normalized chromatin accessibility (left), unspliced pre-mRNA (middle), and spliced mature mRNA (right). Panels show representative genes across cell states: definitive endoderm (**C**), pancreatic progenitor (**D**), early endocrine progenitor (**E**), late endocrine progenitor (**F**), Beta (**G**), Alpha (**H**), Delta (**I**), Exocrine (**J**) and Serotonergic Islet Cells (**K**). Dots represent individual cells and are colored by gene state: primed (red; accessibility increases while expression remains low), coupled-on (orange; both accessibility and expression increase), decoupled (green; accessibility and expression change in opposite directions), and coupled-off (blue; both decrease). Lines show the mean within each gene time bin. Gene time is the inferred temporal ordering for each gene’s accessibility-to-RNA transition.

**Fig. S7 Integrative characterization of data generated using the protocol reported by Balboa et. al.**

**(A)** UMAP (30,465 cells) colored by sequencing modality. **(B)** UMAP colored by differentiation day. **(C)** UMAP colored by annotated cell state. **(D)** Stacked bar graph showing the number of cells across days. **(E)** Dot plot showing gene expression of selected markers. Prolif., proliferative cells; EP, endocrine progenitor; Mesen., mesenchyme; SIC, serotonergic islet cells.

**Fig. S8 Integrative characterization of data generated using the protocol reported by Hua et. al.**

(**A**) UMAP (5,962 cells) colored by sequencing modality. (**B**) UMAP colored by differentiation day. **(C)** UMAP colored by annotated cell state. (**D**) Stacked bar graph showing the number of cells across days. (**E**) Dot plot showing gene expression of selected markers. Prolif., proliferative cells; EP, endocrine progenitor; SIC, serotonergic islet cells.

**Fig. S9 Integrative characterization of data generated using the protocol reported by Hogrebe et. al.**

(**A**) UMAP (124,840 cells) colored by sequencing modality. (**B**) UMAP colored by differentiation day. (**C**) UMAP colored by annotated cell state. (**D**) Stacked bar graph showing the number of cells across days. (**E**) Dot plot showing gene expression of selected markers. DE, definitive endoderm; PGT, primitive gut tube; Prolif., proliferative cells; PP, pancreatic progenitor; EP, endocrine progenitor; SIC, serotonergic islet cells.

**Fig. S10 Integrative characterization of data generated using the protocol reported by Rezania et. al.**

(**A**) UMAP (40,412 cells) colored by sequencing modality. (**B**) UMAP colored by differentiation day. (**C**) UMAP colored by annotated cell state. (**D**) Stacked bar graph showing the number of cells across days. (**E**) Dot plot showing gene expression of selected markers. DE, definitive endoderm; PGT, primitive gut tube; Prolif., proliferative cells; PP, pancreatic progenitor; EP, endocrine progenitor; EXO, exocrine; UE, undefined endoderm progenitor; Mesen., mesenchyme; SIC, serotonergic islet cells.

**Fig. S11 Integrative characterization of data generated using the protocol reported by Veres et. al.**

(**A**) UMAP (41,171 cells)colored by sequencing modality. (**B**) UMAP colored by differentiation day. (**C**) UMAP colored by annotated cell state. (**D**) Stacked bar graph showing the number of cells across days. (**E**) Dot plot showing gene expression of selected markers. DE, definitive endoderm; PGT, primitive gut tube; Prolif., proliferative cells; PP, pancreatic progenitor; EP, endocrine progenitor; EXO, exocrine; UE, undefined endoderm progenitor; Mesen., mesenchyme; SIC, serotonergic islet cells.

**Fig. S12. Integrative characterization of data generated using the protocol reported by Weng et. al.**

(**A**) UMAP (4,635 cells) colored by sequencing modality. (**B**) UMAP colored by differentiation day. (**C**) UMAP colored by annotated cell state. (**D**) Stacked bar graph showing the number of cells across days. (**E**) Dot plot showing gene expression of selected markers. DE, definitive endoderm; PGT, primitive gut tube; Prolif., proliferative cells; PP, pancreatic progenitor; EP, endocrine progenitor; EXO, exocrine; UE, undefined endoderm progenitor; Mesen., mesenchyme; SIC, serotonergic islet cells.

**Fig. S13 Integrative characterization of data generated using the protocol reported by Zhu et. al.**

(**A**) UMAP (153,118 cells) colored by sequencing modality. (**B**) UMAP colored by differentiation day. (**C**) UMAP colored by annotated cell state. (**D**) Stacked bar graph showing the number of cells across days. (**E**) Dot plot showing gene expression of selected markers. DE, definitive endoderm; PGT, primitive gut tube; Prolif., proliferative cells; PP, pancreatic progenitor; EP, endocrine progenitor; EXO, exocrine; UE, undefined endoderm progenitor; Mesen., mesenchyme; SIC, serotonergic islet cells.

**Fig. S14 Cell fate transition probability calculated based on multi-omics single-cell optimal transport.**

Probabilistically matched early to late transitions calculated as individual optimal transport problems for each consecutive stage transition.

**Fig. S15 Additional analyses of early SC-islet differentiation subset and TF perturbations.**

(**A**) UMAP embedding of the digital model subset focused on early pancreatic differentiation (8,684 cells), colored by differentiation protocol. (**B**) Dot plot summarizing expression of developmentally relevant marker genes across cell states during early SC-islet differentiation. Dot size indicates the fraction of cells expressing each gene, and color indicates scaled average expression. (**C**) Feature plots showing normalized expression of key cell state marker genes. (**D**) Top 20 genes ranked by degree centrality in definitive endoderm (left), primitive gut tube (middle), and pancreatic progenitors (right). (**E**) Eight selected transcription factor (TF) knockout (KO) simulations shown as vector fields with perturbation scores (top) and the corresponding feature plots showing normalized TF expression (bottom). Negative perturbation scores (purple) predict blockage of the original developmental flow, whereas positive perturbation scores (green) predict promotion. Normalized TF expression is shown to contextualize predicted KO effects in the relevant cell populations.

**Fig. S16 Integrated single-cell RNA-seq atlas of human fetal pancreas and stem cell-derived pancreatic differentiation.**

(**A**) Dot plot summarizing expression of developmentally relevant marker genes across cell states. Dot size indicates the fraction of cells expressing each gene, and color indicates scaled average expression. (**B**) UMAP of the integrated atlas of early pancreatic differentiation (8,684 cells), colored by dataset. (**C**) UMAP of fetal cells only, colored by post-conceptional week. (**D**) UMAP of stem cell-derived cells only, colored by differentiation day.

**Fig. S17 Comparative differential gene expression analysis of in vitro and in vivo pancreatic development.**

(**A–D**) Volcano plots of differentially expressed genes between stem cell-derived and fetal cells within endocrine progenitors (EP; A), Beta cells (B), Alpha cells (C), and Delta cells (D). Blue and yellow points denote significant genes enriched in stem cell-derived or fetal cells, respectively. Bars above each plot indicate the number of significant genes per origin, and selected genes are labeled. (**E**) Dot plot summarizing expression of selected TFs, comparing stem cell-derived (blue) and fetal (yellow) cells across cell states spanning pancreatic development. Dot size indicates the fraction of cells expressing each gene, and color indicates scaled average expression. PP, pancreatic progenitor; EP, endocrine progenitor

**Fig. S18 Additional analyses of the *in vitro* endocrine and exocrine lineage branching subset and TF perturbations.**

(**A**) UMAP embedding of the digital model subset focused on the endocrine and exocrine lineage branching point (6,726 cells), colored by differentiation protocol. (**B**) Dot plot summarizing expression of developmentally relevant marker genes across cell states within this branching subset. Dot size indicates the fraction of cells expressing each gene, and color indicates scaled average expression. (**C**) Feature plots showing normalized expression of key cell state marker genes. (**D**) Top 20 genes ranked by degree centrality, shown left to right for exocrine (EXO), pancreatic progenitor-exocrine intermediate (PP-EXO), pancreatic progenitor (PP), pancreatic progenitor-early endocrine progenitor intermediate (PP-EEP), and early endocrine progenitor (EEP). (**E**) Eight selected TF knockout (KO) simulations shown as vector fields with perturbation scores (top) and the corresponding feature plots showing normalized TF expression (bottom). Negative perturbation scores (purple) predict blockage of the original developmental flow, whereas positive perturbation scores (green) predict promotion. Normalized TF expression is shown to contextualize predicted KO effects in the relevant cell populations. (**F**) Schematic of doxycycline-inducible CRISPRi-mediated *STAT1* knockdown from days 9-13 during pancreatic progenitor specification. (**G**) qPCR confirming *STAT1* knockdown at day 13. (**H**) qPCR showing that *STAT1* expression returns to baseline by day 26 (no significant difference relative to untreated control). (**I**) qPCR of endocrine markers (left) and exocrine markers (right) at day 26. Data are shown as mean ± s.e.m. relative to untreated control (n = 3 biologically independent samples). Statistical significance was assessed by unpaired two-sided t-test. * P < 0.05.

PP, pancreatic progenitor; PP-EXO, pancreatic progenitor-exocrine intermediate; PP-EEP, pancreatic progenitor-early endocrine progenitor intermediate; EEP, early endocrine progenitor; EXO, exocrine.

**Fig. S19 Pancreatic progenitor-to-Alpha lineage shows dual trajectories towards Alpha cells and reveals regulators of Alpha fate during SC-islet differentiation.**

(**A**) UMAP of the digital model focused on the Alpha lineage (16,122 cells), colored by cell state. (**B**) UMAP colored by differentiation day. (**C**) UMAP colored by differentiation protocol. (**D**) Dot plot summarizing expression of developmentally relevant marker genes across cell states within this subset. Dot size indicates the fraction of cells expressing each gene, and color indicates scaled average expression. (**E**) Feature plots showing normalized expression of key cell state marker genes. (**F**) UMAP colored by gene expression-based pseudotime, with arrows indicating inferred developmental flow. (**G**) Top 15 genes ranked by degree centrality, shown left to right for pancreatic progenitor (PP), endocrine progenitor (EP), early Alpha 1, early Alpha 2, and Alpha cells. (**H–I**) Scatter plots summarizing systematic TF knockout (KO) simulations. Each point represents a TF, positioned by mean perturbation score for PP (x-axis) and early Alpha 1 (y-axis) fates (H), or EP (x-axis) and early Alpha 2 (y-axis) fates (I). Quadrants indicate predicted effects: Q1, promote both fates; Q2, suppress the x-axis fate and promote the y-axis fate; Q3, suppress both fates; Q4, promote the x-axis fate and suppress the y-axis fate. Selected TFs are labeled and color-coded by quadrant (Q1 green, Q2 orange, Q3 red, Q4 blue). (**J**) Four selected transcription factor (TF) KO simulations shown as vector fields with perturbation scores (top) and the corresponding feature plots showing normalized TF expression (bottom). Negative perturbation scores (purple) predict blockage of the original developmental flow, whereas positive perturbation scores (green) predict promotion. Normalized TF expression is shown to contextualize predicted KO effects in the relevant cell populations.

**Fig. S20 Pancreatic progenitor-to-Delta lineage reveals regulators of Delta fate during SC-islet differentiation.**

(**A**) UMAP of the digital model focused on the Delta lineage (7,583 cells), colored by cell state. (**B**) UMAP colored by differentiation day. (**C**) UMAP colored by differentiation protocol. (**D**) Dot plot summarizing expression of developmentally relevant marker genes across cell states within this subset. Dot size indicates the fraction of cells expressing each gene, and color indicates scaled average expression. (**E**) Feature plots showing normalized expression of key cell state marker genes. (**F**) UMAP colored by gene expression-based pseudotime, with arrows indicating inferred developmental flow. (**G**) Top 15 genes ranked by degree centrality, shown left to right for pancreatic progenitor (PP), endocrine progenitor (EP), early Delta, and Delta cells. (**H–I**) Scatter plots summarizing systematic TF knockout (KO) simulations. Each point represents a TF, positioned by mean perturbation score for EP (x-axis) and early Delta (y-axis) fates (H), or early Delta (x-axis) and Delta (y-axis) fates (I). Quadrants indicate predicted effects: Q1, promote both fates; Q2, suppress the x-axis fate and promote the y-axis fate; Q3, suppress both fates; Q4, promote the x-axis fate and suppress the y-axis fate. Selected TFs are labeled and color-coded by quadrant (Q1 green, Q2 orange, Q3 red, Q4 blue). (**J**) Four selected transcription factor (TF) KO simulations shown as vector fields with perturbation scores (top) and the corresponding feature plots showing normalized TF expression (bottom). Negative perturbation scores (purple) predict blockage of the original developmental flow, whereas positive perturbation scores (green) predict promotion. Normalized TF expression is shown to contextualize predicted KO effects in the relevant cell populations.

**Fig. S21. Additional analyses of the *in vitro* beta cell and serotonergic islet cell (SIC) lineage branching subset and TF perturbations.**

(**A**) UMAP embedding of the digital model subset focused on the beta cell and SIC lineage branching point (6,922 cells), colored by differentiation protocol. (**B**) Dot plot summarizing expression of developmentally relevant marker genes across cell states within this branching subset. Dot size indicates the fraction of cells expressing each gene, and color indicates scaled average expression. (**C**) Feature plots showing normalized expression of key cell state marker genes. (**D**) Top 20 genes ranked by degree centrality, shown left to right for early endocrine progenitor (EEP), late endocrine progenitor (LEP), SIC, beta-SIC intermediate progenitors (beta-SIC), and Beta cells. (**E**) Eight selected TF knockout (KO) simulations shown as vector fields with perturbation scores (top) and the corresponding feature plots showing normalized TF expression (bottom). Negative perturbation scores (purple) predict blockage of the original developmental flow, whereas positive perturbation scores (green) predict promotion. Normalized TF expression is shown to contextualize predicted KO effects in the relevant cell populations. EEP, early endocrine progenitor; LEP, late endocrine progenitor; SIC, serotonergic islet cells; Hybrid, population expressing markers of both beta and SIC markers.

## STAR METHODS

### KEY RESOURCES TABLE

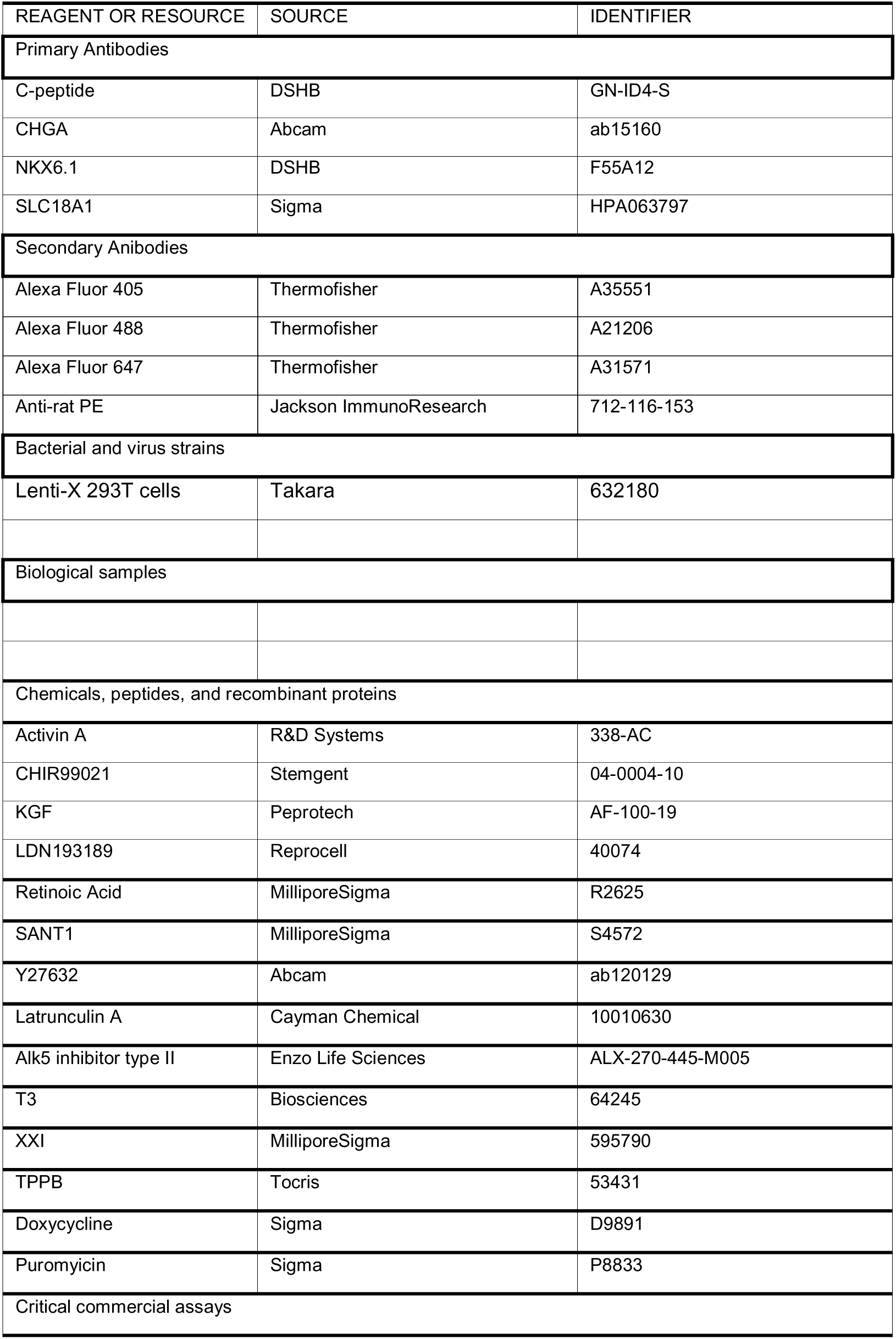
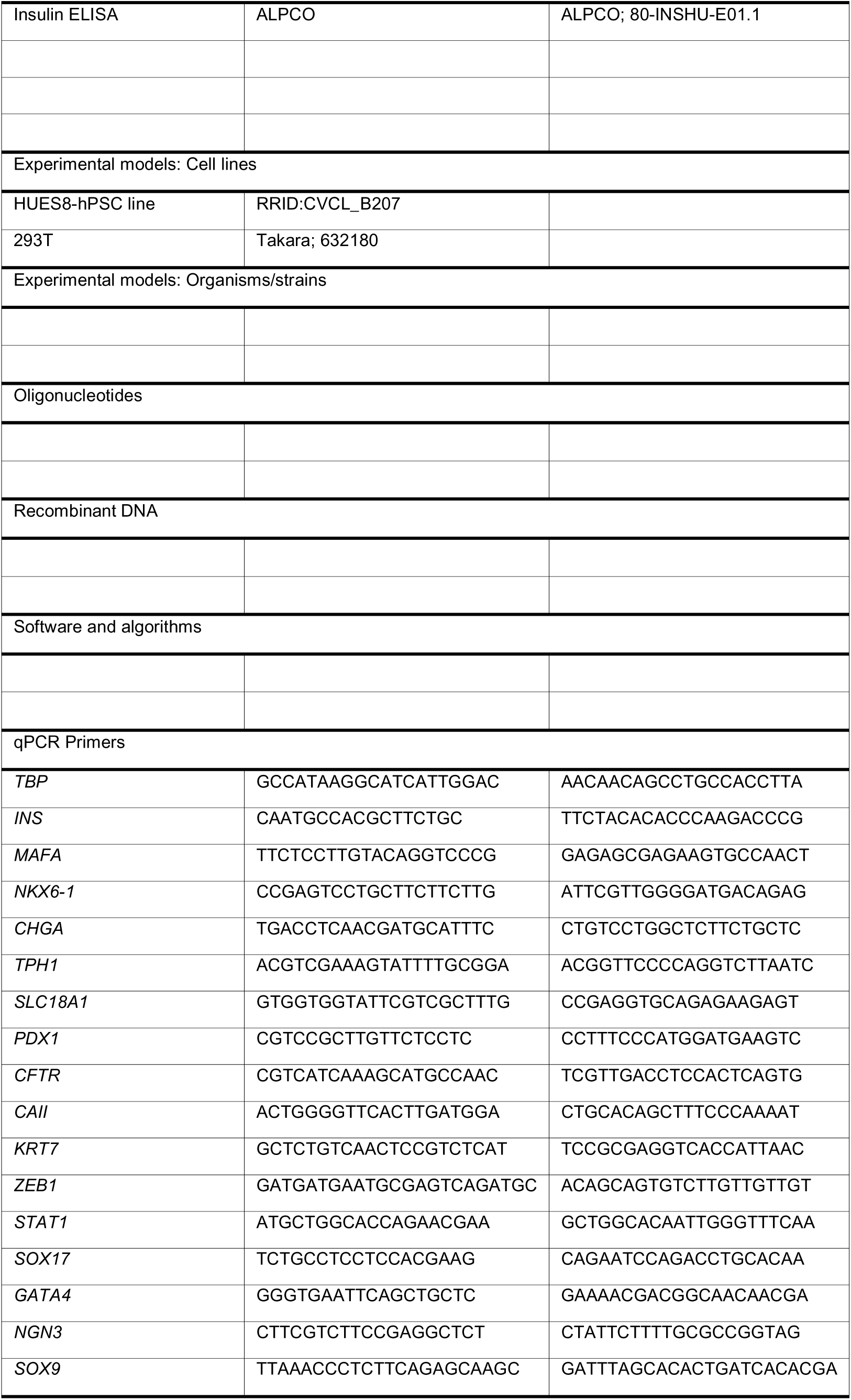

## METHOD DETAILS

### Stem cell culture and differentiation

The Washington University Embryonic Stem Cell Research Oversight Committee approved all work involving human embryonic stem cells (hESCs, Approval number 15-002). HUES8 hESCs (RRID:CVCL_B207), provided by Douglas Melton (Harvard University), were maintained on T75 tissue culture flasks coated with Matrigel (Corning; 356230) in a humidified incubator at 5% CO2 and 37 °C. Cells were passaged approximately every 4□days by first washing with phosphate-buffered saline (PBS) and then incubating with TrypLE (Gibco; 12 604-013) for 10 min or less at 37 °C. Dispersed cells were then mixed with an equal volume of mTeSR1 supplemented with 10 µM Y-27632 (Pepro Tech; 129382310MG) and counted on a Vi Cell XR Cell Viability Analyzer (Beckman Coulter). Cell suspensions were then centrifuged at 300g for 3□min at room temperature (RT). The supernatant was aspirated, and cells were resuspended at a concentration of 1×10^6^□cells/mL and seeded onto Matrigel-coated plates at a density of 0.5×10^5^ cells/cm^2^ for propagation or 5.0×10^5^ cells/cm^2^ for differentiation. Differentiation of hESCs into stem cell-derived islets was performed as described previously^14^, with media formulations and timings outlined in **Table S25**.

### Immunocytochemistry

Samples were fixed with 4% paraformaldehyde for 30□min at RT. Samples were then blocked for 45□min at RT with PBS□+□0.1% Triton-X 100□+□5% donkey serum. The primary antibody was incubated overnight at 4□°C. After washing with PBS, samples were incubated with secondary antibodies for two hours at room temperature. DAPI (4′,6-diamidino-2-phenylindole) was used to stain nuclei. Imaging was performed on a Leica DMI4000 B Inverted Microscope. Antibody details can be found in Table S25.

### Quantitative real-time polymerase Chain Reaction (qPCR)

Cell clusters were collected, washed, and resuspended in an RLT buffer. RNA was extracted from SC-islets using the RNesy Mini Kit (QIAGEN; 74016) and DNase treatment (QIAGEN; 79254). cDNA was then synthesized using the High-Capacity cDNA Reverse Transcriptase Kit (Applied Biosystems; 4368814) and a T100 thermocycler (Bio-Rad). PowerUp SYBR Green Master Mix (Applied Biosystems; A25741) generated real-time PCR reactions on the QuantStudio™ 6 Pro System. Data was analyzed using ΔCt to calculate normalized expression and the ΔΔCt methodology to calculate expression fold change. *TBP* was used as a housekeeping gene for normalization. Primer sequences are available in **Table S25**.

### Static glucose-stimulated insulin secretion assay (sGSIS)

To conduct sGSIS, clusters were placed in a transwell transwell (MilliporeSigma, PIXP01250) and washed with 1□mL of Krebs-Ringer Bicarbonate three times. Then the transwells were transferred to 2□mM glucose for 1 h to equilibrate cells. Next, cells were moved into 2□mM glucose for 1□h followed by 20□mM for 1□hour. Supernatant was collected for both 2□mM and 20 mM. Insulin concentration was measured using enzyme-linked immunosorbent assay (ELISA) kits (ALPCO; 80-INSHU-E01.1), following the manufacturer’s instructions. Hormone secretion values were normalized to cell counts.

### Dynamic glucose-stimulated insulin secretion assay

Dynamic GSIS was performed using the Biorep Technologies Perifusion System (Model No. PER4-115). SC-islets were sandwiched between two layers of hydrated Bio-Gel P-4 polyacrylamide beads (150-4124, Bio-Rad) in 275-μl cell chambers (Peri-Chamber, Biorep Technologies. Samples were then perifused with 2□mM glucose KRB solution for 1 hour at a flow rate of 100□µl min^−1^. After this equilibration period, effluent was collected in 2-min time intervals, switching glucose solutions as follows: 2□mM glucose KRB for 12□min, 20□mM glucose KRB for 24□min and 2 mM glucose KRB for 16□min. The clusters were then lysed with a solution of 10□mM Tris (T6066, MilliporeSigma), 1□mM EDTA (AM9261, Ambion) and 0.2% Triton X (327371000, Acros Organics).DNA was quantified using the Quant-iT Picogreen dsDNA assay kit (P7589, Invitrogen) and was used to normalize insulin values quantified with a human insulin ELISA kit (ALPCO; 80-INSHU-E01.1).

### Transplantation studies

Hyperglycemia was induced in 7-week-old male immunodeficient mice (NOD.Cg-Prkdcscid Il2rgtm1Wjl/SzJ, Jackson Laboratories) by injection with 45 mg/kg streptozotocin (STZ; 1621500, R&D Systems) in 0.9% sterile saline (51-405022.052, Moltox) for 5 consecutive days. Mice were diabetic (blood glucose > 250 mg/dL) 9 days after the final STZ injection. Mice were randomly assigned to experimental groups, anesthetized with isoflurane and injected with approximately 2.25×10^6^ SC-islet cells in the kidney capsule. Mice were monitored for up to 12 weeks. Each week, random and fasting blood glucose levels (4-6 hour fast) were recorded for each mouse. Animal studies were performed in accordance with Washington University International Care and Use Committee (IACUC) regulations (protocol #21-0240).

### Glucose tolerance test (GTT)

To assess glucose tolerance, mice were fasted for 5-6 hours then received an intraperitoneal injection of glucose (3 g/kg body weight) dissolved in 0.9% sterile saline (Moltox, 51-405022.052). Blood glucose levels were measured using a glucometer prior to injection and at four 30-min intervals. C-peptide levels were measured using an ultrasensitive human C-peptide ELISA (Mercodia, 10-1141-01).

### Single-nucleus multiomic sequencing (snMulti-seq)

Samples were processed by first dispersing cells into a single-cell suspension using TrypLE for up to 15 min at 37 °C. Cell number and viability were quantified using a Vi-Cell XR Cell Analyzer (Beckman Coulter). Nuclei were isolated from single-cell suspensions according to the 10x Genomics Multiome ATAC□+□Gene Expression (GEX) protocol (CGOOO338). Briefly, samples were washed with PBS supplemented with 0.04% v/v BSA, lysed with chilled Lysis Buffer for 4□min, washed three times with wash buffer, and then resuspended in 10X nuclei buffer at 3,000–5,000 nuclei/μL. Samples were delivered to the McDonnell Genome Institute at Washington University for library preparation and sequencing. Isolated nuclei were processed using the 10x Chromium instrument, with a target of 7,000–10,000 cells, and the 10x Genomics Single Cell Multiome ATAC□+□Gene Expression v1 kit was used according to the manufacturer’s instructions for library preparations. The NovaSeq 6000 System (Illumina) was used for library sequencing.

### snMulti-seq data processing

snMulti-seq data were processed and aligned to the human reference genome assembly GRCh38 using Cell Ranger ARC (v2.0). A matrix of peaks per cell was constructed for each sample based on ATAC-seq fragments measured per sample across a common peak set generated from all samples. Initial quality control was performed on each sample individually to remove low quality cells based on the total number of RNA molecules detected per cell, total number of ATAC peaks per cell, approximate ratio of mononucleosomal to nucleosome-free fragments (nucleosome signal), and transcriptional start site enrichment (TSSe) score. We removed nuclei with more than 40,000 RNA molecules or ATAC peaks per cell. Nuclei with less than 1,000 RNA molecules or ATAC peaks per cell were also removed. Nuclei with a TSSe score less than 2 were removed. Nuclei with a nucleosome signal greater than 1.5 were removed, except for the Day 26 and Day 33 samples where the filter threshold was set to 5. Final peak calling was performed using Model-based Analysis of ChIP-Seq (MACS2)^80^. The standard Seurat preprocessing workflow was used to normalize, scale, and center RNA data^81^. Linear dimensional reduction was performed by principal component analysis, as implemented in Seurat. Normalization and linear dimensional reduction of ATAC data was performed using term frequency-inverse document frequency (TF-IDF) and singular value decomposition (SVD) functions, respectively, as implemented in Signac^76^. Joint Uniform Manifold Approximation and Projection (UMAP) of RNA and ATAC data was constructed based on a weighted nearest neighbor (WNN) calculated from both RNA and ATAC dimensional reductions.

### snMulti-seq data integration

To integrate snMulti-seq datasets, we first merged data from individual time points during SC-islet differentiation. Next, RNA data was renormalized and scaled then integrated using the streamlined integration method implemented in Seurat v5. We assessed 4 different methods (merging data, CCA, RPCA, and Harmony) using the single-cell integration benchmarking pipeline^82–84^. To integrate ATAC data, we defined a set of anchors using reciprocal LSI projection to create a shared low-dimensional latent space. MACS2 was used to call peaks within the integrated object and joint UMAP visualization was constructed based on a WNN graph of the integrated linear reductions. DNA sequence motif information was added from JASPAR2024 database^85^. ChromVAR was used to determine variations in chromatin accessibility^86^.

### Global chromatin accessibility scoring

The deepTools *computeMatrix* tool was used to calculates scores per TSS±500 base pairs based on hg38v48^87^. The *plotHeatmap* and *plotProfile* tools were used to create profile plots for calculated scores.

### Curation of additional sequencing data

Additional scRNA-seq, snRNA-seq, snATAC-seq, and snMulti-seq datasets were obtained from publicly available repositories (**Supplementary Tables 5 and 11**). Raw FASTQ files were downloaded from the Gene Expression Omnibus (GEO) database (https://www.ncbi.nlm.nih.gov/geo/) using the NCBI Sequencing Read Archive (SRA) toolkit (https://github.com/ncbi/sra-tools). For RNA and ATAC sequencing samples processed using 10x Genomics workflows, data were processed and aligned to human reference genome assembly GRCh38 using Cell Ranger (v7.2.0) and Cell Ranger ATAC (v2.1.0), respectively. For samples processed using inDrops or Drop-seq workflows, aligned raw counts were used for downstream processing and analysis.

### Quality control filtering

Initial quality control was performed on each sample individually to remove low quality cells on a per sample basis. RNA data were filtered based on number of RNA molecules detected per cell, number of genes detected per cell, and percentage of reads mapped to the mitochondrial genome. ATAC data were filtered based on nucleosome banding pattern and transcriptional start site enrichment score.

### Multimodal integration

Multimodal integration of scRNA-seq, snRNA-seq, snATAC-seq, and snMulti-seq datasets was performed using MultiVI ^88^. Briefly, individual scRNA-seq datasets were initially preprocessed to remove low quality cells based on the total number of RNA molecules detected per cell, total number of genes per cell, and percentage of mitochondrial genes. RNA datasets were integrated using Harmony. Next, individual ATAC peak sets were aggregated to ensure that there are common features across all datasets. Low quality cells were removed based on total number of ATAC nucleosome signal, and TSSe score. The combined ATAC data was then integrated using reciprocal LSI projection^84^. Next, the MultiVI model was setup with each modality (RNA, ATAC, or paired) as the main batch annotation, individual datasets as a categorical covariate. The model was trained using paired snMulti-seq data as an anchor within a latent space with 50 dimensions and 3 hidden layers for the encoder and decoder neural networks.

### Diffusion pseudotime

Pseudotime analysis was performed using Monocle3^34^. Specific cell populations representing progenitor lineage divergence were isolated from the integrated Seurat object using the subset function. The resulting subset was converted to a Monocle-compatible cell_data_set object using as.cell_data_set, retaining UMAP embeddings from the original object. Cells were clustered, and developmental trajectories were inferred using cluster_cells with variable settings according to each case. Root cells were manually defined based on known developmental stages using the order_cells function, and pseudotime values were computed and added back to the Seurat object using the AddMetaData function.

### Multiomic stochastic velocity analysis

We used MultiVelo to model the joint dynamics of chromatin accessibility, unspliced RNA, and spliced RNA during *in vitro* differentiation^35^. Prior to analysis, we identified differentially expressed genes using the Wilcoxon rank-sum test, retaining genes with fold change >1.25 and Bonferroni-adjusted *P* < 0.05, and excluding uncharacterized or non-coding transcripts. MultiVelo was then used to infer latent gene time and to model gene-specific regulatory dynamics from paired chromatin accessibility and RNA measurements. Based on the fitted trajectories, cells were assigned, for each gene, to four regulatory gene states: primed, in which chromatin accessibility increases while expression remains low; coupled-on, in which accessibility and expression increase together; decoupled, in which accessibility and expression change in opposite directions; and coupled-off, in which both accessibility and expression decrease. In parallel, we related these MultiVelo-derived dynamics to independently inferred cellular RNA-based pseudotime, calculated as previously described, to examine how chromatin accessibility, unspliced RNA, and spliced RNA changed across differentiation progression.

### Multi-modal trajectory analysis

We reconstructed trajectories from scRNA-seq, snRNA-seq, snATAC-seq, and snMulti-seq data within the digital model using MOSCOT (v0.5.0) to map cells across known time points^89^. Briefly, a cost matrix was generated using the square Euclidian distances within the multimodal latent space generated by MultiVI. Then local PCAs were computed for each pair of time points and solved as individual optimal transport problems per time point pair to probabilistically match early to late transitions into transport matrices. A Markov chain was derived from multiomic transition probability matrices using the *RealTimeKernel* in CellRank (v2.0.7) and cellular dynamics were recovered using the *plot_random_walks* function^90^. Cell state transitions were calculated by aggregating transport matrices using MOSCOT’s cell_transition function. Accuracy of the cell state transition model was determined by root mean square error (RMSE) calculated by:

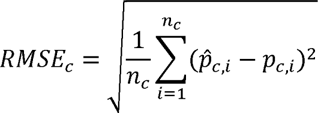

Where *n_c_* 1461 is the number of evaluated stage transitions in which cell state *c* was present in the target stage, *p̂_c,i_* is the predicted proportion of cell state *c* for transition *i*, and *p_ci_*, 1462 is the corresponding observed proportion. Predicted proportions were obtained by applying the transition matrix to the observed proportions in the preceding stage, and observed proportions were calculated from measured target-stage cell counts normalized to sum to 1 across the cell states included in that transition.

### Gene regulatory network inference and transcription factor in silico perturbations

Gene regulatory network (GRN) inference was performed using CellOracle (v0.12.0)^58^ with integrated chromatin accessibility and transcriptomic data. A base GRN was first constructed from the scATAC-seq component by identifying accessible peaks containing candidate transcription factor binding motifs and linking regulatory elements to target genes using co-accessibility and transcription start site annotation. Weak peak-to-gene associations were removed to generate the final prior network.

To construct the Oracle object, this base GRN was combined with the corresponding only scRNA-seq subset of cells from the digital model, cell-state annotations, and Monocle3-derived pseudotime trajectories. Before GRN inference, the top 3,000 highly variable genes were selected using sc.pp.filter_genes_dispersion with the Seurat flavor. After KNN-based imputation, cell-state-specific GRNs were inferred using the get_links function. To reduce spurious interactions, inferred edges were filtered by statistical significance (*P* < 0.001) and ranked by edge strength, retaining the top 2,000 edges for downstream analyses.

*In silico* transcription factor perturbation simulations were then performed across transcription factors represented in the inferred networks to predict their effects on transcriptional state and cell-state dynamics. For each perturbation, CellOracle computed a perturbation score (PS) as the inner product between the pseudotime gradient vector field, representing the direction of natural differentiation, and the simulated perturbation vector field, representing the predicted cell-state transition after transcription factor perturbation. Positive PS values indicate that the simulated perturbation shifts cells in the same direction as the differentiation trajectory, whereas negative PS values indicate a shift opposing differentiation. To summarize lineage-specific effects, we calculated the mean PS within each cell state.

### Comparative analysis in vivo and in vitro pancreatic development

To assess similarities and differences between *in vivo* and *in vitro* pancreatic development, we integrated scRNA-seq datasets from both sources (**Table S11)** after excluding mitochondrial and ribosomal genes. Cells were curated using canonical marker expression and assigned to corresponding pancreatic cell states. Differential expression analysis was then performed within each matched cell state to compare *in vivo* and *in vitro* cells using the Wilcoxon rank-sum test. Transcriptional differences were visualized with volcano plots using a fold-change threshold of 1.5 and an adjusted *P*-value threshold of 1 × 10^-100^. To identify biological programs distinguishing *in vivo* and *in vitro* cells, gene set enrichment analysis was performed on the resulting differentially expressed genes.

### Gene set enrichment analysis

Gene set enrichment analysis was performed using Enrichr-KG^91^. Enrichment was evaluated using the Gene Ontology Biological Process 2021 library^92^. Enrichr computes term enrichment using Fisher’s exact test and reports a z-score reflecting deviation from the expected rank; terms were ranked using Enrichr’s combined score, defined as *c=-In(p)✕z*, where is the Fisher exact test p-value

### Generation of lentiviral constructs

Primer sequences targeting *STAT1* and *ZEB1* were ordered from Millipore Sigma (**Table S25**). To assemble the plasmid, the primers were resuspended in ultrapure water to a concentration of 100□μM. Forward and reverse primers (1□μL each), T4 ligation buffer (NEB, B0202A), T4 Polynucleotide Kinase (NEB, M0201S), and Ultrapure water were added to a PCR tube and placed in the thermocycler for phosphorylation and annealing reaction. The resulting gRNA oligonucleotide duplex was inserted into a Lentiguide Puro Backbone (Addgene plasmid #73797) by combining Rapid Ligase buffer (Enzymatics, B1010L), Fast Digest Esp31 (Thermo, FD0454), dithiothreitol (Promega, PRP1171), bovine serum albumin (NEB, B9000S), and T7 DNA ligase (Enzymatics, L6020L). This master mix was placed in the thermocycler for the Golden Gate assembly reaction to generate plasmids. Plasmids containing the short hairpin RNA (shRNA) sequences for *ZEB1* were purchased as glycerol stocks (MilliporeSigma, TRCN0000017565).

Plasmid DNA was isolated using the QIAprep Miniprep kit (Qiagen, 27115) and then transformed into One Shot Stbl3 Chemically Competent *Escherichia coli* (Invitrogen, C737303). Single colonies were picked after 18-24 h and further expanded cultured. DNA extraction was performed using the Qiagen Maxi plus kit (Qiagen, 12981). Lentiviral particles were generated using the Lenti-X 293T cells (Takara, 632180) cultured in 10 cm plates (Falcon, 353003) with Dulbecco’s modified Eagle medium, 10% heat-inactivated fetal bovine serum (MilliporeSigma, F4135), and 0.01 mM sodium pyruvate (Corning, 25-000-CL). Once confluent, the Lenti-X 293T cells were transfected with 6 μg of plasmid DNA, 4.5 μg of psPAX2 (Addgene, 12260), 1.5 μg pMD2.G (Addgene, 12259), and 48 μL of Polyethylenimine Max (Polysciences, 24765-2). Media was changed after 16 h. Virus containing supernatant was collected at 96 h post-transfection, concentrated using the Lenti-X concentrator (Takara, 631232), and tittered using Lenti-X GoStix Plus (Takara, 631280).

### Generation of CRISPR inactivation cell line

To generate a doxycycline-inducible CRISPR inactivation (CRISPRi), the pT077 KRAB-dCas9-IRES-EGFP AAVS1 targeting plasmid (Addgene # 137879) was used^93^. For targeted integration, AAVS1 TALEN L (Addgene #59025) and AAVS1 TALEN R (Addgene #59026) plasmids were co-transfected with the donor plasmid. All plasmids were amplified in E. coli, purified using a Maxi-prep kit (Qiagen), and resuspended in sterile, nuclease-free water to a final concentration of 1 µg/µL. For transfection, 1×10^6^ HUES8 cells were resuspended in 100 µL of supplemented P3 Nucleofector™ solution (from P3 Primary Cell 4D-Nucleofector® X Kit, S; Lonza). The cell suspension was mixed with 5 µg of total plasmid DNA, consisting of a mixture of the pT076 donor and the two TALEN plasmids. Cells were transfected using the Amaxa™ 4D-Nucleofector™ X Unit (Lonza) with program CM-113. Immediately following nucleofection, cells were plated onto Matrigel-coated plates in mTeSR™ Plus medium supplemented with CloneR™ (STEMCELL Technologies) to enhance single-cell cloning efficiency. Selection with Geneticin (50 µg/mL, Fisher Scientific; cat. no. 10131035) was initiated 48 hours post-transfection. After 10-14 days, emerging colonies were treated with 2 µg/mL doxycycline to induce KRAB-dCas9-IRES-EGFP expression. GFP-positive colonies were manually isolated, expanded, and cryopreserved for subsequent experiments.

### Gene Knockdown via lentiviral transduction

For CRISPRi gene knockdown, HUES8 cells stably expressing doxycycline-inducible dCas9-KRAB were transfected with lentivirus contained guide RNA for either *STAT1* or *ZEB1* at multiplicity of infection (MOI) of 10. Transduced cells were expanded in media supplemented with 0.4 µg/mL puromycin for 3 passages and subsequently seeded for differentiation following the standard SC-islet generation protocol. Doxycycline (10□μg/mL) was added to induce dCas9-KRAB expression during from days 9 to 12.

For gene knockdown via shRNA, lentivirus containing sequences targeting *ZEB1* was added to culture media on days 15 and 16 of the differentiation protocol at a MOI of 10. As a control, cells were treated with lentivirus containing shRNA targeting a GFP gene sequence.

### Ruxolitinib treatment

Ruxolitinib was added to the culture media at a final concentratrion 20□μM from days 9 to 13 of the SC-islet differentiation protocol. Media changes were performed according to the standard differentiation schedule. All other steps in the protocol remained unchanged. Additional details regarding Ruxolitinib are provided in Table S25.

### Flow cytometry

Cells were single-cell dispersed by adding 0.2 ml TrypLE/cm^−2^ for 10 min at 37 °C. Cells were washed with PBS, centrifuged, and fixed by resuspending them in 4% paraformaldehyde at room temperature for 30 minutes. After another PBS wash, samples were then blocked for 45□min at room temperature with PBS□+□0.1% Triton-X 100□+□5% donkey serum. Primary antibodies were incubated overnight at 4□°C. The secondary antibodies were incubated the next day for two hours at room temperature. Antibody details and dilutions can be found in Table S25. Cells were washed twice with PBS and then filtered before being run on the Cytek Northern Lights flow cytometer using SpectroFlow. FlowJo v10.10 (Becton, Dickinson, and Company) was used for analysis.

### Hormone content assessment

Hormone content was measured by cells, rinsing with PBS, and incubating in an acid–ethanol solution for 48 hr at −20°C. Samples were neutralized with 1 M TRIS buffer (Millipore Sigma; T6066), and hormone content was quantified using ELISA kits for human insulin (ALPCO; 80-INSHU-E01.1) and serotonin (ALPCO; 17-SERHU-E01-FST). ELISAs were performed according to the manufacturer’s instructions, and results were normalized to cell counts.

## QUANTIFICATION AND STATISTICAL ANALYSIS

The statistical analyses were performed using GraphPad Prism 10.2.2 software (GraphPad Software, San Diego, CA). Statistical analysis for qPCR, flow cytometry, and hormone content assays was performing using multiple unpaired t-tests without assuming consistent standard deviation, no correction for multiple comparisons, and 0.05 set as the threshold for statistical significance. Two-way analysis of variance (ANOVA) was used to calculate statistical significance for sGSIS assays. All values were expressed as mean ± s.e.m. and corresponding P values are reported in individual figure captions. Sample sizes were not predetermined using statistical methods. Experiments were conducted without randomization, and investigators were aware of group allocation during both experimentation and data analysis.

## ADDITIONAL RESOURCES

Interactive online tool of the digital model of stem cell-derived islet differentiation: https://www.synbioelab.com/islettwin

